# The amino acid transporter CG1139 is required for retrograde transport and fast recovery of gut enterocytes after *Serratia marcescens* intestinal infection

**DOI:** 10.1101/2021.10.29.466403

**Authors:** Catherine Socha, Inês S. Pais, Kwang-Zin Lee, Matthieu Lestradet, Dominique Ferrandon

## Abstract

The intestinal tract is constantly exposed to microbes. Severe infections can arise following the ingestion of pathogenic microbes from contaminating food or water sources. The host directly fights off ingested pathogens with resistance mechanisms, the immune response, or withstands and repairs the damages inflicted either by virulence factors or the host immune effectors, through tolerance/resilience mechanisms. In a previous study, we reported the existence in *Drosophila melanogaster* of a novel evolutionarily conserved resilience mechanism to intestinal infections with a hemolysin-positive *Serratia marcescens* strain (*Sm*Db11), the purge of the apical cytoplasm of enterocytes. The epithelium becomes very thin and recovers rapidly, regaining its normal thickness within several hours. Here, we found that this recovery of gut enterocyte morphology is based on the host internal reserves and not on ingested food. Indeed, we observed a retrograde transport of amino acids from the host hemolymph to the enterocytes. We have identified several amino acid transporters required for recovery and we focused on the SLC36 family transporter CG1139. *CG1139* is required for the retrograde transport of amino acids. RNA sequencing revealed that genes involved in the positive regulation of growth were observed in wild-type but not *CG1139* mutant guts, in which the expression of *Myc* and genes involved in Insulin signaling is down-regulated. Functional analysis revealed that *Myc* is also required for the recovery of the thick gut epithelium after infection. Altogether, our results show the importance of an amino acid transporter in the fast regrowth of the enterocytes upon infection. Unexpectedly, we found that this transporter acts non cell-autonomously and can regulate the transcription of other genes, suggesting a signaling function of CG1139 that therefore appears to act as a transceptor.

## Introduction

The intestinal tract is one of the largest interfaces between the host body and the external environment, harboring a diverse microbial community that is mostly beneficial to the host (Flint, Scott, Louis, & Duncan, 2012). However, the proliferation of existing opportunistic pathogens as well as the exposure to food-borne pathogens can severely compromise host health (Ryu et al., 2008). Resistance and resilience mechanisms in the host have been selected during evolution that ensure the homeostasis of the epithelium and limit damages inflicted by pathogens (Ferrandon, 2013). Whereas resistance mechanisms are responsible for decreasing the pathogen burden, resilience mechanisms are responsible for enduring the presence of pathogens. *Drosophila melanogaster* is a powerful model organism to study such mechanisms, especially due to the power of its sophisticated genetic tools. The regulation of intestinal stem cells (ISCs) is an integrated response required for gut homeostasis and for the replacement of the gut enterocytes that are lost over time (Bonfini, Liu, & Buchon, 2016).

Amino acids have also been shown to be important in the outcome of an infection (Zhu, Deng, & Yin, 2018). Amino acid metabolism affects the host physiology and may serve as an energy source for the cell. In mammals, 59 amino acid transporters have been identified and categorized in 12 different Solute Carrier (SLC) families (Thwaites & Anderson, 2011) such as the Proton-coupled Amino acid Transporters (SLC36 or PATs), the Sodium-coupled Neutral Amino acid Transporters (SLC38 or SNATs), the cationic (SLC7 or CATs) and the heterodimeric (SLC7/SCL3 or HATs) amino acid transporters. In *Drosophila*, among the average 14 000 genes, 603 are predicted to encode putative transporter proteins, which correspond to 4% of the genome (Featherstone, 2011). Amino acid transporters have redundant functions, as different transporters can carry the same amino acid. These characteristics allow adaptation to various environmental conditions.

*Serratia marcescens* (*Sm*) is a Gram-negative opportunistic human pathogen that secretes several virulence factors, such as proteases and pore-forming toxins. Hemocyte-mediated phagocytosis in the body cavity is an effective response against the ingested *Sm* bacteria that manage to cross the intestinal barrier. When injected directly in the hemolymph, *Sm* can resist the systemic immune response essentially because of the O-antigen of its LPS (C. L. Kurz et al., 2003; Nehme et al., 2007). Several genes required for susceptibility or for resistance against ingested *Sm* have been identified before in a genome-wide screen. The JAK-STAT pathway was shown to be required for host defence by regulating the proliferation of ISCs (Cronin et al., 2009), a result independently obtained by other investigators (Apidianakis, Pitsouli, Perrimon, & Rahme, 2009; N Buchon, Broderick, Chakrabarti, & Lemaitre, 2009; Jiang et al., 2009).

We have previously reported the cytoplasmic purge of enterocytes, a very fast response of the gut epithelium to hemolysin expressing *Serratia marcescens* (*Sm*Db11) (Lee et al., 2016). After ingestion, the enterocytes of the midgut extrude a considerable portion of their cytoplasm through an aperture that is formed in the apical part of the cell. This happens at very early stages, around one hour upon infection. Consequently, the cells become very thin and flat; enterocytes of the R2 region lose their dome-shaped apical domains typical of this portion of the midgut. Strikingly, the cytoplasmic purge prevents damages to enterocytes as there is no increased enterocyte cell death nor compensatory stem cell proliferation. Moreover, the same cells are able to recover their normal shape and size within the first 20 hours post infection. Damaged organelles such as mitochondria and also likely toxins and invading bacteria are actively extruded out of the exposed enterocytes. We have previously identified *Cyclin J* (*CycJ*) as being required for the recovery of the cell size and shape (Lee et al., 2016). However, it remains unclear how the host can recover from a very thin epithelium that lost a massive part of its cytoplasm and associated organelles and reconstitute the normal cell shape and size. Here, we explore this question and show that a retrograde transport of metabolites occurs from the hemolymph to the enterocytes during the recovery of the epithelium. We found that the amino acid transporter CG1139 localizes basally in the labyrinth region of the enterocytes, a zone of intensive exchange between the enterocyte and the hemolymph, and is required for the retrograde transport of a modified amino-acid. Moreover, CG1139 is required for the fast regrowth of the epithelium, but a recovery is still possible in its absence, albeit slower, suggesting a possible redundancy with other amino acid transporters also shown to be involved in the regrowth of thinned enterocytes. Unexpectedly for a transmembrane protein, we found CG1139 to act non-cell autonomously and we suggest that it may be involved in cell to cell signaling and possibly inter-organ communication.

## Results

### A retrograde transport from the hemolymph to the gut takes place upon infection

When flies are orally infected with *S. marcescens* that express hemolysin (*Sm*Db11), midgut enterocytes expel their apical cytoplasm, which can be observed as large cytoplasmic extrusions forming in the ectoperitrophic space. The cells subsequently experience a considerable decrease in their thickness and volume, as well as a lowered microvilli density and size (Lee et al., 2016). As a consequence of this cytoplasmic purge, these cells lose a fraction of their components and organelles, such as lipids and mitochondria (Lee et al., 2016). Strikingly, the recovery to the normal shape and volume occurs relatively rapidly, being completely restored around 16h to 24h after infection. We started by investigating the origin of the reserves that are required for the intestinal epithelium recovery. We usually infect the flies with *Sm*Db11 in a 50mM sucrose solution containing 10% of Lysogeny broth medium (LB). LB is a nutritive medium containing peptides and vitamins that sustains bacterial growth. To analyze a possible role of LB in the recovery and growth of the gut enterocytes after infection in an anterograde manner, *a.k.a*, direct absorption of nutrients by the intestinal epithelium, we replaced LB by PBS and analyzed if the flies recovered to the same extent as the control infected flies fed sucrose solution with 10% LB. Three hours post-infection the cytoplasmic purge occurred to the same extent in both conditions leading to a very thin intestinal epithelium of the gut (**Figure 1A**). At 16h post-infection, the gut enterocytes were able to recover equally well under both conditions (**Figure 1A**). We also replaced LB by a mix of essential and non-essential amino acids in the infection solution as a further positive control and obtained similar results (**Figure 1A**). These results suggest that the presence of LB in the infection solution is not required for the recovery of the gut enterocytes. Because flies were still feeding on sucrose during infection, we decided to analyze if sucrose itself could have some role in the recovery. For this purpose, we exposed the flies to *Sm*Db11 re-suspended in a PBS1x solution for 3h in the absence of any sucrose or LB; after that period of exposure to bacteria, we flipped the flies to sterile PBS1x, to H2O or to sucrose with LB as a control for 13h. Surprisingly, almost all the flies in both conditions were able to restore a thick epithelium in the absence of any external nutrients (**Figure 1B, 1C**, **Supplementary Figure 1A**). These results suggest that upon infection, the fast recovery of the gut epithelial cells from the cytoplasmic purge is not dependent on newly acquired nutrients, but rather on metabolites stored in the insect body.

**Figure 1.**
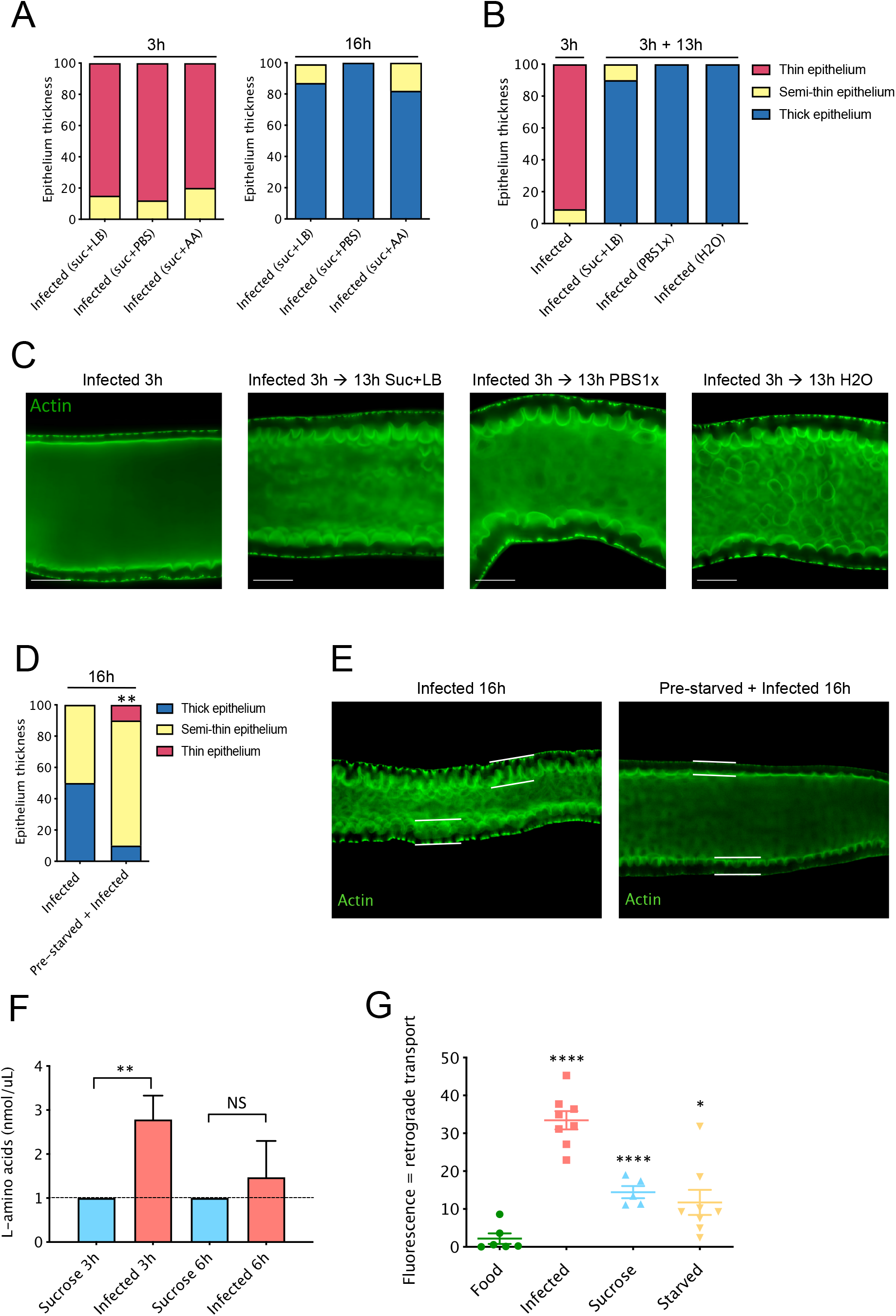
Recovery of enterocytes after infection entails retrograde transport from body reserves. **(A)** Flies were infected in a sucrose solution containing 10% LB medium, PBS1x or a mix of essential and non-essential amino-acids (AA). Thickness of gut epithelium was accessed at 3h and 16h after *Sm*Db11 infection. All conditions recover to a thick epithelium 16h post infection. **(B)** Flies were infected for 3h with *Sm*Db11 suspended in PBS1x solution, and then transferred to sucrose with 10% of LB medium solution, PBS1x or to H2O. Thickness of gut epithelium was accessed at 3h after infection and at 13h after transferring the flies to the second solution. Guts were stained with phalloidin and representative pictures of gut enterocytes from each condition are shown in **(C)**. **(D)** Flies fed in standard food or flies starved for 24h were infected with *Sm*Db11 and thickness of gut epithelium was accessed at 16h post infection. 16h post infection the percentage of midguts with thick epithelia is lower when flies are pre-starved. Representative pictures of gut enterocytes are shown in **(E)**. **(F)** Quantification of free amino-acids in the hemolymph of flies 3h and 6h after sucrose exposure or *Sm*Db11 infection. The level of free amino-acids increases in the hemolymph at 6h (One-way ANOVA, *p* < 0.001). **(G)** Click-it AHA kit was used to assess a retrograde transport from hemolymph to the intestine. Flies were injected with 50 μM of AHA and placed on different conditions: regular food, infected with *Sm*Db11, sucrose 50 mM or water (starved). Midguts were dissected 6h later, and stained with the alkyne probe to assess the fluorescence corresponding to the incorporated amino-acid. Detailed scheme in **Supplementary figure 1B**. Fluorescence was calculated by measuring green (Alexa 488) intensity with the Image J software. The increase in fluorescence was considered as read-out for the level of retrograde transport from the midgut. Infected flies presented the higher level of retrograde transported (One-way ANOVA; *=p<0,05; ****=p<0,0001). Green = Actin. Qualitative quantification of intestines according to their epithelial thickness : thin (red), semithin (yellow) or thick (blue).

To explore this possibility, we decided to starve flies for 24h in H2O to deplete their internal metabolic stores and then to observe how well they would recover from the cytoplasmic purge. Only 10% of the flies were able to recover to a normal epithelium, while the others 80% and 10% displayed respectively a semi-thin and thin epithelium (**Figure 1D, E**). These results show that when flies are pre-starved before the infection, the gut epithelium cannot recover to the same extent as nonstarved flies 16h post-infection.

We reasoned that after infection a retrograde transport of metabolites would be occurring from the rest of the body to the gut to sustain the fast recovery of the epithelium to its normal thickness. We first measured the level of free amino acids in the hemolymph after bacterial infection. At 3h post-infection, we observed an increase of free amino acids in the hemolymph of flies. This increase was transient since it was no longer detected 6h post-infection (**Figure 1F**). This observation led us to ask whether amino acids in the hemolymph might be transported to the gut in a retrograde manner during the recovery process. To answer this question, we used the Click-iT AHA (L-azidohomoalanine) technique, which employs a modified amino acid that is an analog of methionine recognized by the translation machinery and incorporated into newly synthetized proteins. After reacting AHA with an alkyne fluorophore probe, the incorporated analog can be detected by fluorescence in tissues in which translation is taking place (**Supplementary Figure 1B**). We injected AHA in the hemocoel of flies. Next, we exposed the injected flies to either standard fly food, *Sm*Db11, sucrose or to water (starved flies) for 6h. Flies placed on food presented very low or absent fluorescence in the gut in contrast to infected flies, which presented a large increase in the midgut epithelium fluorescence 6h post-infection (**Figure 1G**). Interestingly, flies that were placed in sucrose or starved in water also displayed an increase of the midgut epithelium fluorescence (**Figure 1G**). These results show that the modified injected amino acid is being taken up from the hemolymph and incorporated into the enterocytes during the recovery phase of cytoplasmic extrusion or upon amino-acid starvation. Taken together, these experiments suggest the occurrence of a retrograde transport to the midgut epithelium in response to infection or starvation.

### The amino acid transporter CG1139 is required for the fast recovery from the gut epithelium after the cytoplasmic purge

Because there is an increase in free amino acids in the hemolymph and a retrograde transport to the gut enterocytes upon infection, we next asked if gut amino acid transporters are involved in epithelium recovery upon infection. We screened 28 putative amino acids transporters expressed in the *Drosophila* midgut by knocking down their gene expression in enterocytes using the Gal4/UAS system with a specific driver (NP1-Gal4, Gal80^ts^=NP) and analyzed how the corresponding midguts recover from infection. Interestingly, the knockdown of several amino acid transporter genes in the gut resulted in an impaired recovery of the gut epithelium after the cytoplasmic purge (**Supplementary Figure 2A**), suggesting their involvement in this process. Remarkably, one of the amino acid transporters hit that presented the strongest phenotype with more than 50% of the guts with an epithelium remaining thin, *CG1139*, was the sole transporter gene found to be up-regulated 6h post-infection in an RNA sequencing experiment that was previously performed in our laboratory (**Figure 2A**). We validated the RNA sequencing results by RT-qPCR and we observed a more than 10-fold up-regulation of *CG1139* transcripts at 9h post-infection (**Figure 2B**). We also analyzed the expression of *CG1139* upon infection with *Sm*21C4, a *S. marcescens* mutant for hemolysin that does not cause an accentuated cytoplasmic purge (Lee et al., 2016). Although *CG1139* was still somewhat induced, it was not as strongly expressed as after *Sm*Db11 challenge (**Figure 2B**).

**Figure 2.**
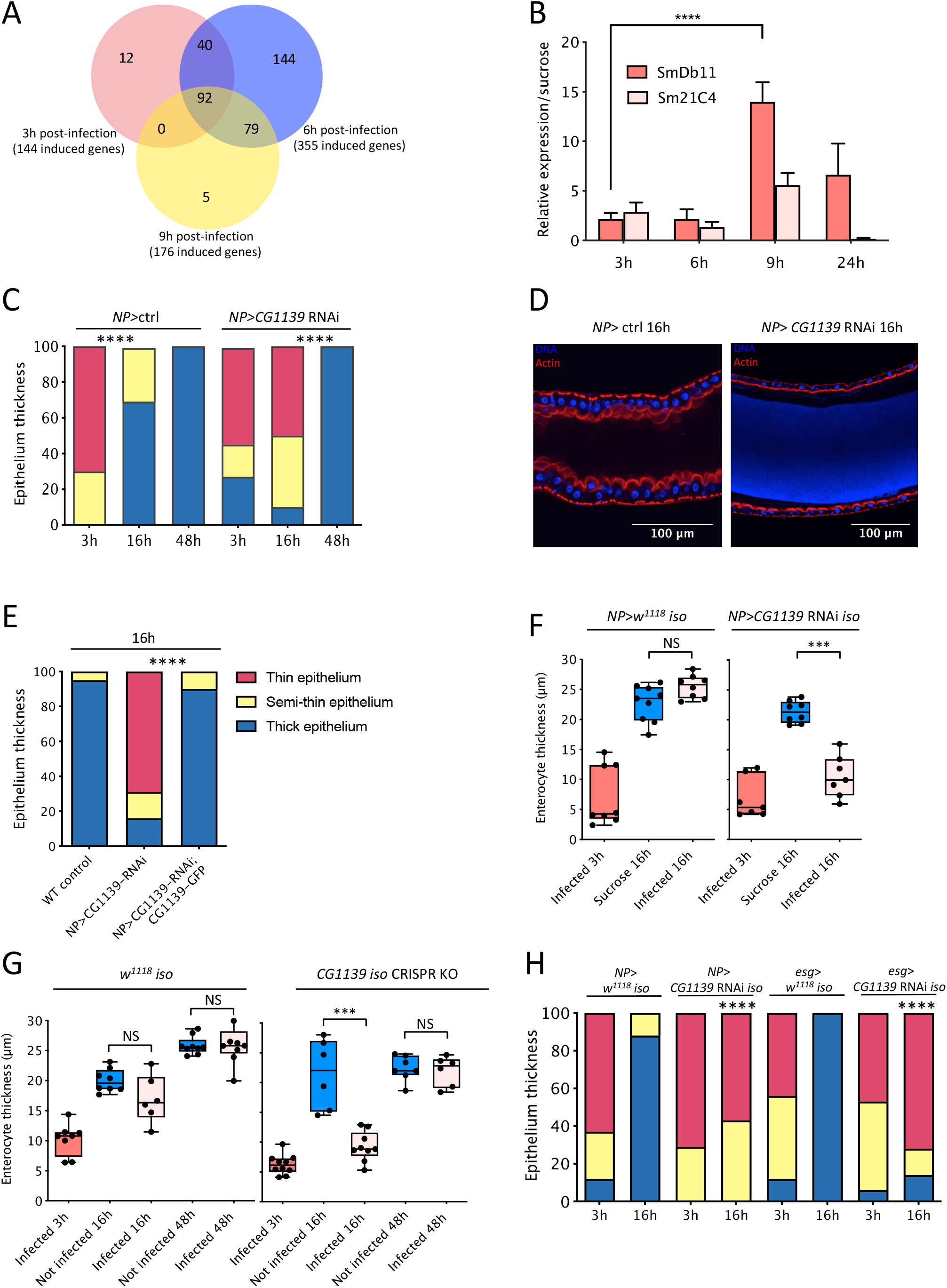
Fast recovery of gut epithelial thickness requires the CG1139 amino acid transporter. **(A)** Genes induced in the midgut during infection with *Sm*Db11 identified by RNA-sequencing. RNA was extracted from 10 midguts in triplicates. 144, 355 and 176 genes are induced 3, 6 and 9h post-infection in the intestine, respectively, and 92 genes are induced throughout the infection. **(B)** *CG1139* expression measured by RT-qPCR. RNA was extracted from 10 midguts, in triplicates. Bars represent expression of infected flies relative to sucrose controls. *CG1139* was induced in the midgut during the recovery, 9h post-infection with *Sm*Db11 (One-way ANOVA, ****=*p* <0,0001) **(C)** Control flies (*NP>ctrl*) and *NP>CG1139*-RNAi were infected with *Sm*Db11, midguts were dissected at 3h, 16h, and 48h post infection and the thickness of the gut epithelium was accessed. Midguts from *NP>CG1139*-RNAi flies are thinner than the control 16h post-infection. **(D)** Representative midguts from control flies and *NP>CG1139*-RNAi flies 16h post-infection. Blue: DNA stained with DAPI; Red: actin stained with phalloidin. **(E)** Epithelium thickness 16h after *Sm*Db11 infection in *NP>*ctrl, *NP>CG1139*-RNAi and in the rescued line *NP>CG1139-* RNAi;UAS-CG1139-GFP. The rescue of *CG1139* rescues the delay on epithelium recovery. **(F)** NP> *w^1118^ iso* and *NP> CG1139* RNAi *iso* were exposed to sucrose or infected with *Sm*Db11 and enterocytes thickness was measured using ImageJ software. Each dot represents the average of 10 independent measures in one midgut. *NP> w^1118^ iso* recover the normal size of the enterocytes thickness compared to sucrose, but enterocytes from *NP> CG1139* RNAi iso remain thin 16h post infection (*lmer*, *** = *p* <0.001). **(G)** *w^1118^ iso* and a Knock-out CRISPR mutant for *CG1139* (*CG1139 iso* CRISPR KO) were exposed to sucrose (Not infected) or infected with *Sm*Db11 and enterocytes thickness was measured as in **(F)**. *CG1139* mutant has a delay in recovery at 16h but recovers the normal enterocyte thickness 48h post infection similar to sucrose condition (*lmer*, *** = *p* <0.001). **(H)** Flies where *CG1139* was knocked down in the enterocytes (*NP*> *CG1139* RNAi iso) or in the ISCs (*esg> CG1139* RNAi *iso*) and the respective controls (*NP> w^1118^ iso* and *esg > w^1118^ iso*) were infected with *Sm*Db11 and midguts were dissected 3h and 16h post infection to assess the epithelium thickness. Both *NP> CG1139* RNAi *iso* and *esg> CG1139* RNAi *iso* present a delay in recovery, with higher percentage of midguts with thin epithelium 16h post infection compared to the respective controls. Quantification of intestines according to their epithelial thickness: thin (red), semithin (yellow) or thick (blue).

*CG1139* encodes an amino acid transporter similar to a mammalian PAT (Proton-assisted SLC36 Amino acid Transporter) (**Supplementary Figure 2B**) and it has previously been shown in *D. melanogaster* to be involved in cell growth (Goberdhan, Meredith, Boyd, & Wilson, 2005). The possibility that this transporter might regulate the recovery phase that follows the cytoplasmic purge prompted us to investigate further the role of this specific transporter in enterocytes.

Having validated the efficiency of *NP>CG1139* RNAi (**Supplementary Figure 3A**), we next confirmed the results from the mini-screen of amino acid transporters by analyzing the gut epithelium recovery of *NP>CG1139* RNAi flies 16h and 48h post-infection. Both *NP>CG1139* RNAi and control flies showed a cytoplasmic purge at 3h, with thin epithelium of the gut enterocytes. After 16h, we confirmed that almost all the control flies had recovered a normal gut epithelium thickness. In contrast, only 10% of *NP>CG1139* RNAi flies had restored the enterocytes back to their normal size, with 50% of the flies still presenting a very thin epithelium of the gut (**Figure 2C, D**). However, 48h post-infection *NP>CG1139* RNAi flies had recovered completely, suggesting that the absence of this transporter causes a delay in the recovery. We obtained similar results using an independent RNAi line, and we also performed a rescue experiment with the overexpression of a transgene encoding a *CG1139-GFP* fusion in the *NP>CG1139* RNAi background. The expression of a wild-type copy of *CG1139* was sufficient to reinstate the recovery by 16h even though a fusion protein was used (**Supplementary Figure 3B**, **Figure 2E**). To limit possible effects of the genetic background of *CG1139* RNAi, we isogenized this line into the *w^1118^ iso* background and measured more precisely the enterocytes thickness. Overall, we obtained similar results to those obtained with the non-isogenized lines reported so far. In sucrose, both NP>*iso* and NP> CG1139 RNAi *iso* presented enterocytes with a thickness around 22μm. At 3h post-infection, when the cytoplasmic purge had occurred, the enterocytes had remarkably lost some 15μm thickness (median of 7μm) (**Figure 2F**). At 16h post-infection, while control flies had recovered their size (median of 25.5μm), enterocytes from NP>*CG1139* RNAi *iso* remained thin (median of 10.5μm), not different from *Sm*Db11 at 3h (lm, *p* = 0.1023).

In addition to RNAi, we used a CRISPR mutant for *CG1139* to confirm the delayed recovery of the gut observed with RNAi lines (**Supplementary Figure 3C**). After infection, there is no up-regulation of *CG1139* in the mutant, similar to *NP>CG1139* RNAi (**Supplementary Figure 3D**). We confirmed that the mutant also presented a delay in the recovery after infection, with the gut enterocytes remaining thin at 16h but recovering after 48h, similar to *CG1139* knockdown in the enterocytes (**Figure 2G**). All together, these results show that *CG1139* is an amino acid transporter induced in the gut upon infection and required in the enterocytes for the fast recovery of the gut epithelium after the cytoplasmic purge.

High-Throughput expression data from Flybase shows that *CG1139* is expressed in the adult midgut, but it is also very highly expressed in the Malpighian tubules. In addition, data from Flygut-*seq* shows that *CG1139* is also expressed in the visceral muscles surrounding the gut. Therefore, we decided to knockdown *CG1139* in tissues and cell types other than enterocytes to observe if the role in recovery would be restricted to this intestinal epithelium cell type. We knocked down its expression ubiquitously (*ubi-Gal4-Gal80^ts^*), in the fat body (*yolk*-*Gal4*), in the Malpighian tubules (*uro*-*Gal4*), in the visceral muscles (*how*-*Gal4*), in the stem cells (*esg*-*Gal4*) and in the enteroendocrine cells (*pros*-*Gal4*). When *CG1139* was knocked down ubiquitously (*ubi>CG1139* RNAi), the flies presented a similarly thin epithelium at 16h post-infection as *NP>CG1139* RNAi flies (**Supplementary Figure 4A**). None of the other flies displayed an impaired recovery when *CG1139* was knocked down in the corresponding tissues (**Supplementary Figure 4A-C**), except for the *esg*-*Gal4* where *CG1139* was downregulated in epithelial progenitor cells (**Figure 2H**, **Supplementary Figure 4D**). More than 70% of *esg*-*Gal4*>*CG1139* RNAi flies displayed a thin epithelium 16h post infection (**Figure 2H**). These results suggest that either *CG1139* is also required in the progenitor cells of the gut epithelium during recovery or that during differentiation the new enterocytes generated during the RNAi induction period that lasts between 5 to 7 days lack the transporter as well.

To visualize the tissues and the cell types in which *CG1139* may be expressed, we used as reporter a *UAS-GFP* transgene crossed to a *CG1139* CRISPR Knock-in mutant we have generated in which the entire CDS region of *CG1139* has been deleted and replaced by Gal4 coding sequences (**Supplementary Figure 5A**). Interestingly, although we observed the expression of *CG1139* in some enterocytes and mostly in the posterior midgut region, we also found that cells with a shape characteristic of progenitor cells express *CG1139*, either in control and infection conditions (**Supplementary Figure 5B**). This observation suggests that CG1139 may also be important at the level of the progenitor cells and explain the delayed recovery obtained using *esg*-*Gal4* driver. Of note, the knock-in line may have deleted some regulatory elements present in the coding regions or introns and may therefore only partially reflect the endogenous expression of *CG1139*.

The cytoplasmic purge has previously been shown to be involved in restricting the number of bacteria that cross the epithelium into the hemolymph (Lee et al., 2016). Here, we asked if the knockdown of *CG1139* in the gut and the associated delay in the recovery would impact the fitness of the host upon infection. We did not observe any difference in fly locomotion or survival upon infection (**Supplementary Figure 6A, B**). However, 16h post-infection when NP>*CG1139* RNAi flies still had not fully recovered from the cytoplasmic purge, the loads of *Sm*Db11in these flies were 10-fold higher in the crop and in the midgut compared to control flies (**Supplementary Figure 6C**). To analyze if this increased bacterial burden resulted from an increased feeding behavior or from constipation caused by *Sm*Db11 ingestion, we monitored the feeding rate of flies as well as their defecation rate. We did not observe any striking difference for these behaviors upon *Sm*Db11 exposure (**Supplementary Figure 6D-F**), suggesting that the higher bacterial loads are related to the delayed recovery.

### CG1139 localizes to the basal part of the gut enterocytes and is required for the retrograde transport

We have shown that a retrograde transport of amino acids is occurring from the hemolymph to the gut during the phase of recovery after the cytoplasmic purge. We hypothesized that the amino acid transporter CG1139 might play a role in the retrograde transport, and that it would be required for the fast recovery of the enterocytes upon infection. *CG1139* has previously been shown to be localized at the cell surface and on the late endosomal and lysosomal surfaces in *Drosophila* S2 cells, as well as in the larval fat body. We started by determining the subcellular localization of CG1139 in the midgut of flies by ectopically expressing in enterocytes the functionally active CG1139-GFP fusion. We detected the green fluorescence in the basal part of enterocytes close to the visceral muscles (**Figure 3A**) in the basal labyrinth region of the cell that allow intensive exchanges of enterocytes with the hemolymph (Shanbhag & Tripathi, 2009). This basal localization was maintained during infection (**Figure 3A**), suggesting that CG1139 is likely involved in the amino acid transport from the hemolymph to the gut.

**Figure 3.**
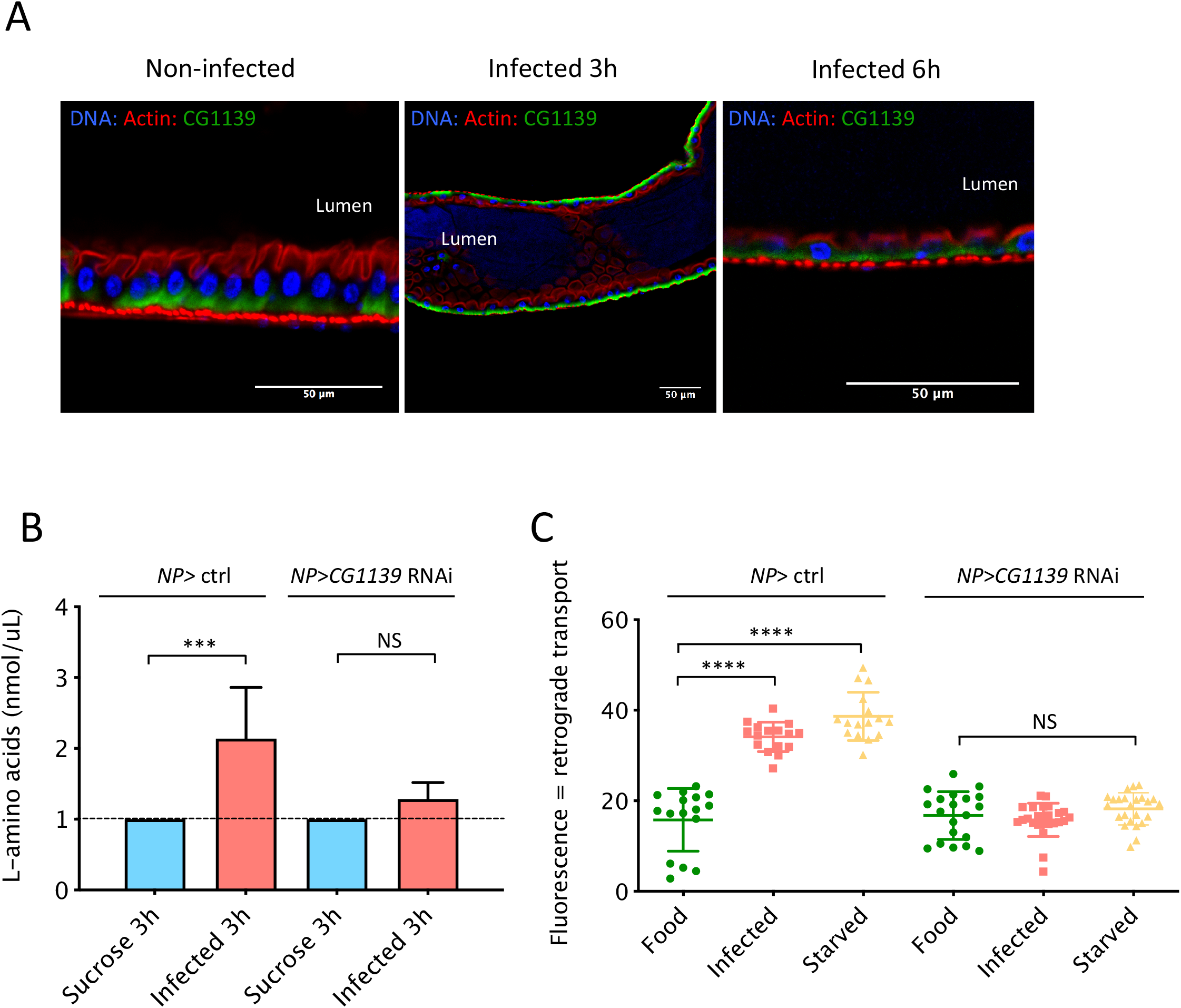
CG1139 localizes to the basal side of the enterocytes in the midgut epithelium and is required for the retrograde transport. **(A)** Confocal pictures of dissected midguts from *NP>UAS-CG1139*-GFP flies exposed to sucrose (Non-infected), or exposed to *Sm*Db11 for 3h or 6h. Blue: DNA; Red: actin; Green: CG1139 GFP. **(B)** Quantification of free amino-acids in the hemolymph of *NP>* ctrl and *NP> CG1139* RNAi 3h after sucrose exposure or *Sm*Db11 infection. The level of free amino-acids increases in the hemolymph of *NP>* ctrl flies but not in *NP> CG1139* RNAi flies (*One-way ANOVA*, *** = *p* < 0.001). **(C)** Click-it AHA kit was used to assess a retrograde transport from hemolymph to the intestine as described in **Figure 1**. There is an increase of retrograde transport to the midgut in *NP>* ctrl flies after infection or starvation but not in *NP> CG1139* RNAi. (*One-way ANOVA*, **** = *p* < 0.0001).

We next asked if the transient increase in the level of free amino acids in the hemolymph 3h post-infection would be more pronounced in *NP*>*CG1139* RNAi flies that would reflect their possible accumulation resulting from an impaired importation into the gut epithelium. Unexpectedly, the reverse phenomenon was observed with NP>*CG1139* RNAi flies not presenting any increase of free amino acids in the hemolymph 3h post-infection, in contrast to the control flies. We next performed the Click-iT AHA technique as described above to analyze if *CG1139* would have a role in the retrograde transport after infection. We observed again the retrograde transport to the gut after *Sm*Db11 infection and after starvation in the control flies (**Figure 3C**). Interestingly, we did not observe the occurrence of a retrograde transport after infection or starvation in *NP*>*CG1139* RNAi flies (**Figure 3C**), showing that *CG1139* is required for the retrograde transport of amino acids into the intestinal epithelium. Altogether, these results suggest that *CG1139* is required for the increase of free amino acids in the hemolymph 3h post-infection and for their retrograde transport to the gut epithelium. Paradoxically, the Click-iT technique is based on a modified methionine that is not expected to be translocated by CG1139, which has been shown to transport alanine, proline, and glycine. These results suggest that CG1139 may act indirectly to support the retrograde transport to the gut epithelium.

Therefore, we decided to determine if *CG1139* acts cell-autonomously during the recovery as would be expected for a transporter. Here, we used the escargot Flip Out line (esgF/O) and performed a clonal analysis as previously described (Jiang et al., 2009). The esgF/O flies were crossed with *CG1139* RNAi (*CG1139* F/O) transgenic flies to generate clones of enterocytes expressing both GFP and RNAi against *CG1139* in the offspring. We reasoned that during recovery, two scenarios could take place (**Figure 4A**): i) both wild type enterocytes and those expressing *CG1139* RNAi/GFP would present a similar recovery at 16h post-infection (either growing up to normal size or remaining thin), suggesting a non-cell autonomous role of *CG1139*; ii) only enterocytes expressing *CG1139* RNAi/GFP would remain thin at 16h, suggesting a cell autonomous role of *CG1139.* We observed that the first scenario took place, whereby all the enterocytes from the gut recovered to the normal size at 16h post-infection (**Figure 4B**), whether in the *CG1139*-deficient clones or in wild-type enterocytes. The size of the clones did not influence the results, whether midguts comprised small size clones made-up of a few cells scattered throughout the epithelium or large-size clones of 15-20 cells expressing *CG1139* RNAi.

**Figure 4.**
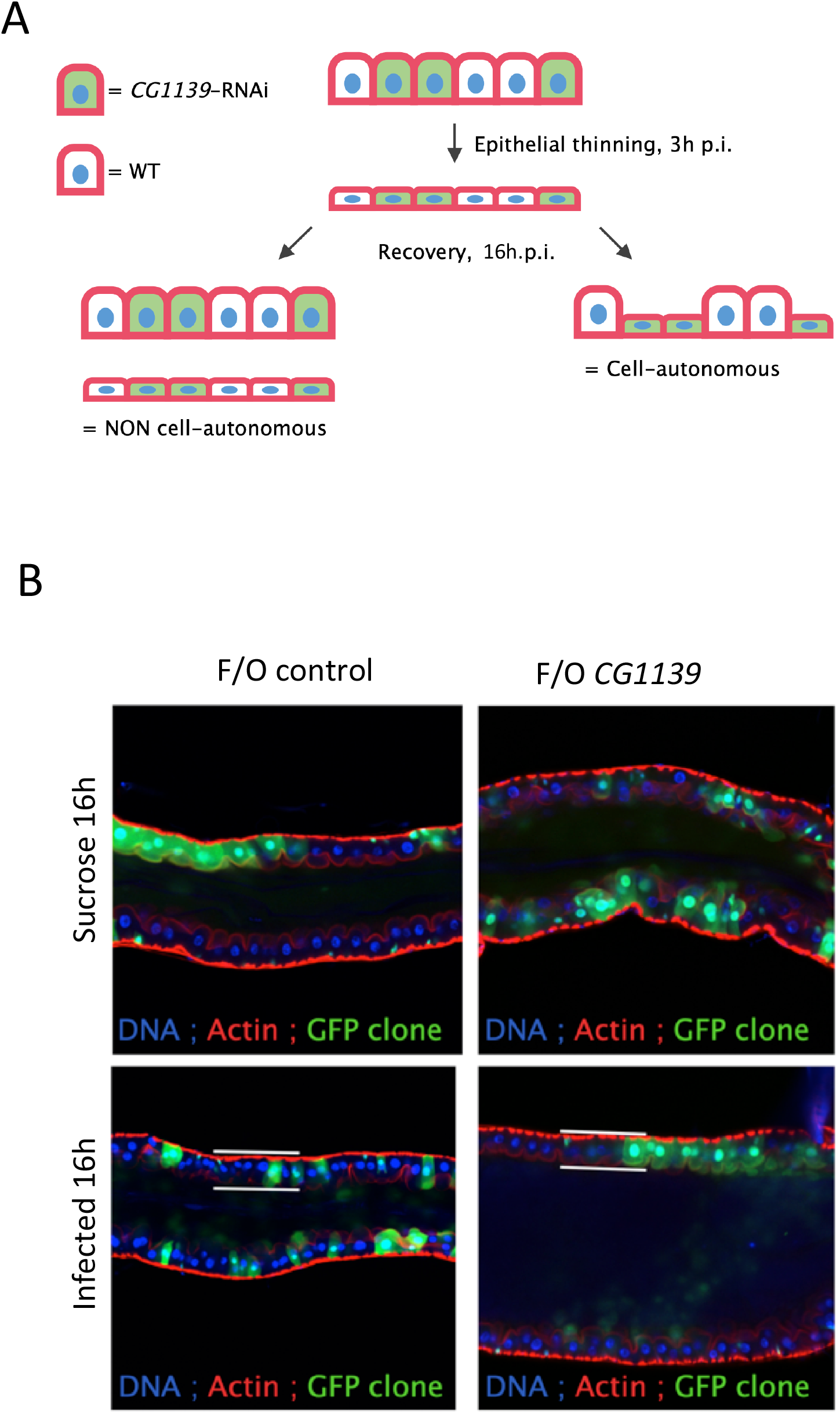
*CG1139* is required non-cell autonomously for the recovery of the intestinal epithelium after *Sm*Db11 infection. Clonal analyses were performed using the escargot Flp Out (*esg*F/O) system. **(A)** *w; esgGal4tubGal80ts UAS-GFP*; *UAS-flp Act>CD2>Gal4* flies were crossed to *UAS-CG1139*-RNAi flies to generate clones of enterocytes expressing both GFP and RNAi against *CG1139* (F/O *CG1139*). Left scenario: both RNAi and wild type enterocyte recover or stay thin 16h post-infection, suggesting a non cell-autonomous mechanism. Right scenario: RNAi enterocytes remain thin and wild-type enterocytes recover, suggesting a cell-autonomous mechanism of *CG1139*. **(B)** Confocal pictures of dissected midguts from control F/O or *CG1139* F/O flies. Blue: DNA; Red: Actin; Green: control GFP clone or *CG1139*-RNAi GFP clone. In F/O control and F/O *CG1139* flies, both RNAi and wild type enterocytes showed the same recovery rate 16h post-infection, suggesting a non cell-autonomous mechanism.

These results show that *CG1139* acts non-cell autonomously, confirming the indirect role of *CG1139* in the recovery of the enterocytes after the cytoplasmic purge triggered by exposure to *Sm*Db11. We conclude that CG1139 does not function solely as a regular amino-acid transporter and ensures additional functions in the control of metabolic fluxes between the rest of the organism and the gut epithelium.

### The transcription factor Myc is required for the fast recovery of the gut enterocytes upon infection

To understand how CG1139 functions to ensure the fast recovery, we performed an RNA sequencing analysis in midguts of control flies or flies for which *CG1139* was knocked down in the enterocytes. We compared genes differentially expressed at 3h, 8h, and 16h upon *Sm*Db11 infection. We performed a principal component analysis (PCA) and we observed that there was a clear-cut difference on the PCA2 axis between noninfected and infected *NP> CG1139* RNAi flies, especially for the 8h and 16h time-points. This difference was less pronounced between uninfected and infected wild-type control samples (**Figure 5A**). We also observed that there were consistently more genes differentially expressed in *NP> CG1139* RNAi than in the wild-type control samples, when comparing the infected samples to the corresponding sucrose controls (**Figure 5B**). To analyze which genes would be differentially regulated upon infection between the wild-type and *NP> CG1139* RNAi, we performed a Gene Set Enrichment Analysis (GSEA) for all the time-points comparing control and mutant knock-down flies to each other. There were several gene sets that were significantly up-regulated in the wild-type compared with the knocked down flies. We focused on the time-points of 8h and 16h, the period during which the intestinal epithelium recovers its initial shape and thickness. Strikingly, at 8h we found an enrichment for genes known to be involved in the positive regulation of organ grown, being up-regulated in the wild-type and not in the *NP> CG1139* RNAi flies. In addition, the gene set of lamellipodium assembly was also enriched and up-regulated in the wild-type and not in the *NP> CG1139* RNAi (**Supplementary Figure 7A**). This suggests that there are genes involved in growth, likely required to the increase thickness of the enterocytes, which were not expressed when we knocked down *CG1139*. Moreover, genes involved in lamellipodium assembly may be involved in actin cytoskeleton, and may be required to close the pore formed during the cytoplasmic purge or to re-shape the enterocyte membrane. Alternatively, they might contribute to the formation of microvilli, which are shortened and lost to some extent upon exposure to *Sm*Db11 hemolysin (Lee et al., 2016). Genes from this category, as well as genes involved in actin nucleation were also up-regulated in the wild-type at 16h (**Supplementary Figure 7B**). Actin nucleation is the first step in the assembly of the actin filaments, and this may have a similar role described above for the lamellipodium assembly related genes.

**Figure 5.**
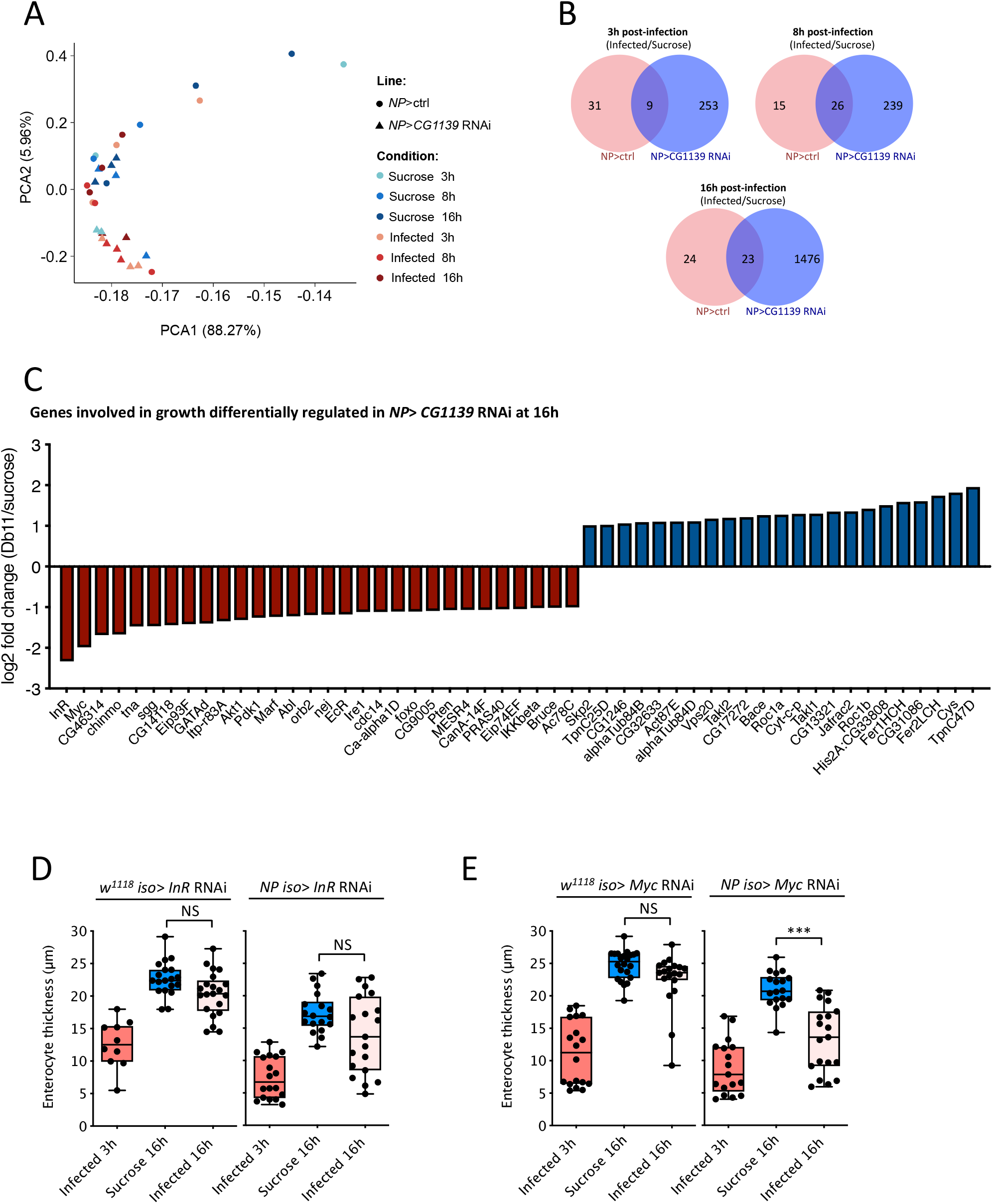
The transcription factor *Myc* is required for the recovery of the gut enterocytes after infection. *NP>* ctrl and *NP>* CG1139 RNAi were exposed to sucrose or infected with *Sm*Db11 for 3h, 8h and 16h. RNA sequencing was performed in dissected midguts. **(A)** Principal component analyses for gene expression of different samples. **(B)** Venn diagram for each time-point comparing genes differentially expressed in *NP>* ctrl and *NP> CG1139* RNAi after infection, relative to sucrose. **(C)** Genes involved in growth that are up- or down-regulated in *CG1139* RNAi 16h after infection (relative to sucrose). Insulin related genes and the growth transcription factor *Myc* are down-regulated in *NP*>*CG1139* RNAi. **(D)** *InR* and **(E)** *Myc* were knocked down in the gut enterocytes (*NPiso*> *Inr* RNAi and *NPiso* > *Myc* RNAi) and compared with the respective controls (*w^1118^ iso*> *Inr* RNAi and *w^1118^ iso*> *Myc* RNAi) for the recovery of the gut enterocytes thickness 16h upon infection or exposure to sucrose. All the flies present a thin epithelium at 3h upon infection. While the thickness of the gut enterocytes in *NPiso*> *Inr* RNAi 16h post infection are not significantly different from the sucrose control, NP*iso*> *Myc* RNAi are not able to recover the normal enterocyte thickness compared to the sucrose control (*lmer*, *** = *p*<0.001).

To analyze potential candidate genes involved in growth that would be differently regulated between the wild-type and the *CG1139* RNAi, we extracted the expression levels of growth related genes from the RNAseq data set and compared infected flies versus their corresponding sucrose control (**Figure 5C**, **Supplementary Figure 8**). Interestingly, we noticed that at 16h when the enterocytes are still thin in *NP> CG1139* RNAi, these flies downregulated several genes involved with the Insulin signaling pathway, such as *Akt1* and *InR* (**Figure 5C**). Because CG1139 was reported before to interact with TOR, we tested if the activation of TOR would be different in these flies. We used immunohistology stainings of midguts with antibodies against the phosphorylated form of 4EBP (P-4EBP) as read-out. At 16h after infection, TOR activity was indeed decreased in *CG1139* RNAi flies compared to the wild-type controls, in keeping with the possibility that CG1139 contributes to the regulation of TOR activity after infection (**Supplementary Figure 9**).

We also directly tested the role of the Insulin Receptor gene *InR*, which was the most downregulated gene in CG1139 RNAi. We knocked down *InR* in the gut enterocytes using the NP-Gal4 driver. In two independent experiments, we observed either that there was a recovery of the knocked-down *InR* flies or alternatively no recovery of the enterocyte thickness, resulting in a bimodal distribution when the two data sets are pooled (**Figure 5D**). We also focused on the second most down-regulated gene found in *CG1139* RNAi flies, which is the gene encoding the MYC transcription factor. This gene is homologous to vertebrate *Myc* proto-oncogenes, is involved in cell growth and it was previously reported to be required for the crypt morphogenesis in the intestine in mice (Bettess et al., 2005). We confirmed that the expression of *Myc* is significantly downregulated after the ingestion of *Sm*Db11 in *NP-Gal4* > *CG1139* RNAi compared to control flies at 16h (**Supplementary Figure 8B,***lmer*, *p*-value = 0.0116). We decided to further investigate the function of *Myc* by testing whether knocking down this gene in the gut enterocytes would impair their recovery. While the control flies *w^1118^ iso*> *myc* RNAi recovered the normal thickness at 16h post infection, NP *iso*> *myc* RNAi flies presented a thinner epithelium and were unable to recover the normal size of the gut enterocytes (**Figure 5E**). These results suggest that *Myc* expression is downregulated in the *CG1139* RNAi flies and that MYC is required for the regrowth of the enterocytes after infection.

Altogether these results indicate that CG1139 seems to regulate the expression of several other genes, such as genes involved in growth, Insulin signaling, and TOR pathway activity. One of these indirect target genes, *Myc*, is also required for the intestinal epithelium recovery after infection. Thus, the relatively reduced expression of *Myc* in *NP>CG1139* RNAi midguts may contribute to their delayed recovery of epithelium thickness.

## Discussion

Ever since the discovery of intestinal stem cells in *Drosophila* (Micchelli & Perrimon, 2006; Ohlstein & Spradling, 2006) and the interest in intestinal infections (Liehl, Blight, Vodovar, Boccard, & Lemaitre, 2006; Nehme et al., 2007; Ji-Hwan Ryu et al., 2006), there has been a renewed interest in studying the homeostasis of the intestinal epithelium in normal and pathogenic conditions (Nicolas Buchon, Silverman, & Cherry, 2014; Lemaitre & Miguel-Aliaga, 2012; De Navascués et al. 2012; O’Brien et al, 2011). A recent study underlined the importance of the diet in determining the size of the *Drosophila* gut and emphasized that the adaptation to novel environmental conditions involves not only the proliferation of ISCs but also on enterocyte growth (Bonfini et al., 2021). As TOR signaling contributes to this adaptation, enterocyte growth is implicitly thought to be a cell-autonomous event. Indeed, the mechanisms behind the control of enterocyte growth and regulation of size have gain interest on the field (Xiang et al., 2017; Wen et al., 2017; Tamamouna and Panagi et al, 2020) The enterocyte cytoplasmic purge provides a unique setting in which the regrowth of enterocytes following the extrusion of their apical cytoplasm is uncoupled from ISC proliferation (Lee et al., 2016). Here, we document that the regrowth of enterocytes depends solely on an inversion of metabolic fluxes from the organism to the gut. We demonstrate that the retrograde transport of amino acids involves the *CG1139* transporter, which is required for the fast regrowth of enterocytes. Unexpectedly, the function of this transporter is shown genetically to be noncell autonomous.

### Retrograde transport of metabolites

During nutrient uptake, absorption of oligopeptides and amino acids is made through an influx in the apical part of the enterocytes. Once inside the cell, oligopeptides are hydrolyzed by peptidases into their constituent amino acids. While 10% of these amino acids are used for intracellular protein synthesis, the rest is exported to the internal milieu through an efflux in the basal part of the membrane. The reverse, an influx of amino acids from the blood to the epithelial cells takes place in crypt enterocytes, which do not absorb nutrients from the crypt lumen (Boron & Boulpaep, 2009). The cytoplasmic purge results in a massive loss of apical cytoplasm for the cells, with a loss of up to 90% of their total volume in the most severe cases. Enterocytes need to recover their shapes fast in order to maintain their normal function. Indeed, recovery happens within the first 16h to 20h post infection. For this process, it is likely advantageous for the host not to rely on potentially contaminated external food sources, but rather to use internal reserves to reconstitute the organelles and missing cytoplasm and to grow back the enterocytes to their normal size. In starvation conditions, it may be important to maintain enterocyte fitness by the import of amino-acids so as to be able to rapidly digest any food found by the fly. During recovery, we show that the uptake of amino acids in the hemolymph by enterocytes is preceded by a transient increase in their concentration. The retrograde transport of a modified methionine to the gut enterocytes is not observed in the absence of the amino acid transporter *CG1139*. As recovery is delayed in *CG1139* mutant guts, it is likely that the uptake of other amino-acids is also impaired. These amino acids are most likely employed to synthetize new proteins in the cells.

A retrograde transport of amino acids was also observed in flies that were subjected to starvation and sucrose. One explanation may be that under both conditions there is the perception of amino acid starvation, and the organism may need to re-allocate amino acids from the rest of the body to the enterocytes. The gut epithelium of flies exposed to full starvation or amino-acid starvation (sucrose) does not become very thin as when flies as subjected to *Sm*Db11. This suggests that the retrograde transport of amino acids does not seem to be directly correlated to the size of enterocytes.

### Basal localization of CG1139 in the membrane of the gut enterocytes

We were surprised to detect CG1139 on the baso-lateral side of the midgut epithelium. Indeed, PATs are usually localized to the apical side of the epithelium, where the pH is lower, thereby transporting metabolites originating from the degradation of dietary nutrients. PAT1 was detected apically in Caco2 cells and in the rat intestine (Anderson et al., 2004; Z. Chen et al., 2003) as well as in *Aedes aegypti* midgut (Evans, Aimanova, & Gill, 2009). However, an electrophysiology study in the larvae revealed the presence of a H+ -ATPase in the basal part of the intestinal epithelium, which establishes a low pH gradient peaking at 4.34 near the basal labyrinth (Shanbhag & Tripathi, 2005). The localization of *CG1139* in the basal part of the enterocytes is consistent with its putative role in the retrograde transport since CG1139 is a transporter of the SLC36 family of symporters that imports amino-acids concomitantly with a H^+^ proton.

An important question is that of the exact subcellular location of CG1139. PATs such as CG1139 have been described on intracellular compartments such as lysosomes and on the cell surface in *Drosophila* (Ogmundsdottir et al., 2012). CG1139 may localize both on the cell membrane in the basal labyrinth and on lysosomes where it can interact with the TOR pathway machinery. It will be interesting to characterize its subcellular distribution or redistribution using endosomal/lysosomal markers.

### The role of CG1139 in the enterocyte size recovery and its redundancy

Although we observed that *CG1139* is required for the rapid recovery of the normal size of cells after infection, the enterocytes still recover their normal size 48h later, in the RNAi knock down flies as well as in the presumably null CRISPR KO mutant. This suggests that other genes are involved in this process, and that this is an integrated response not solely relying on a single gene. The fact that other amino acid transporters were required for the recovery of the gut epithelium leads us to think that they can be acting together in this process and their role may be redundant. While CG1139 belongs to the SLC36 family of proton-coupled amino acid transporters, most of the amino acid transporters that were shown to be required for the recovery belong to the SLC7 family, especially to the cationic amino acid transporters. These transporters are responsible for the major influx of cationic amino acids and important for nitric oxide synthesis by delivering L-arginine for nitric oxide synthase (Fotiadis, Kanai, & Palacín, 2013). Some of the transporters of the SLC7 family have been shown to be involved in growth. *Minidisc* is required for the growth of imaginal discs in a non-cell autonomous manner (Martin, Hersperger, Simcox, & Shearn, 2000) and the transporter it encodes has been shown to transport leucine into Insulin Producing Cells (IPCs) in the *Drosophila* brain, thereby allowing the secretion of insulin-like peptides (Manière, Ziegler, Geillon, Featherstone, & Grosjean, 2016). Slimfast was shown to regulate organismal growth in the fat body by interacting with TSC/TOR signaling and interfering with PI3-kinase signaling in peripheral tissues. Mutation of *slimfast* induces systemic growth defects in larvae and leads to smaller adults, similarly to phenotypes observed upon nutrient deprivation or altered TOR signaling (Colombani et al., 2003).

Of note, when we performed the Click-it experiment we established that *CG1139* is required for the transport of the labeled methionine analogue from the hemolymph to the gut. However, CG1139 has been shown to transport alanine, proline and glycine in xenopus eggs (Goberdhan et al., 2005). Although we cannot formally exclude the possibility that CG1139 can also transport methionine *in vivo*, one possibility is that one of the other amino acid transporters identified here to be required for the recovery is transporting the labeled methionine. Although *CG1139* was the only amino acid transporter found to be up-regulated during the recovery phase, it will be interesting to further analyze the gene expression of the other amino acid transporters at later time-points. Also, it will be worth determining whether these other amino acid transporters are also localized basally on the cell and required for the retrograde transport of amino acids.

### CG1139 and its role in signaling

Several lines of evidence suggest that the role of *CG1139* goes beyond its simple function as an amino acid transporter. In addition to its non-cell autonomous function for the recovery of the gut enterocytes thickness after infection, it was required for the apparition of the peak of amino acids in the hemolymph that is observed in the hemolymph three hours post-infection. This suggests an early role for CG1139 during infection, or that CG1139 establishes a difference in the host physiology *a priori*, since the knock down is performed for a few days at the adult stage prior to the experimental infection. It will be interesting to determine the origin of these amino acids. They could be generated though the degradation of proteins in muscles or in the fat body, but could also directly originate from hemolymph proteins. Indeed, lipoproteins found in the hemolymph are large proteins, potentially generating a consequent amount of metabolites when degraded.

In addition, *CG1139* was also required to modulate the expression of different genes in our RNA sequencing analysis. Of note, genes annotated to be involved in growth in the Gene Ontology analysis were up-regulated in the wild-type during the recovery period. In contrast, some genes involved in Insulin signaling and growth, were more down-regulated in the *CG1139* knocked down flies. *Myc* belongs to a family of proto-oncogenes and was previously shown in *Drosophila melanogaster* to be required for larval growth and endoreplication (Johnston, Prober, Edgar, Eisenman, & Gallant, 1999; Pierce et al., 2004). MYC proteins have also been shown to be involved in cell proliferation and differentiation. In mammals, *c-Myc* was shown to be required in the intestinal mucosa for the formation of the correct number of crypts in the small intestine (Bettess et al., 2005). Further rescue experiments will be necessary to assess if *Myc* is sufficient to rescue the delayed phenotype in *CG1139* and if it acts cell autonomously. Also, it will be interesting to explore if the absence of *Myc* is responsible for a delayed recovery or a complete lack of recovery, also at 48 hours and later.

*CG1139* has previously been shown to interact with TOR pathway and to synergize with its growth promoting effects (Goberdhan et al., 2005; Ogmundsdottir et al., 2012). We also observed lower TOR activity when *CG1139* was knocked down in the enterocytes, either after sucrose or *Sm*Db11 exposure. Indeed, feeding flies with sucrose solution represents an amino-acid starvation condition, which can explain this phenotype. mTORC1 increases ribosome numbers and activity, which promotes the translation of mRNA and stimulates cell growth and proliferation. The main amino acid activator of mTORC1 is leucine, although it can also be activated by glutamine, arginine and serine. CG1139 is not known to transport either of these amino acids. To account for this discrepancy, CG1139 was recently suggested by other group to have a signaling function and to act as a transceptor (Fan & Goberdhan, 2018). Transceptors have been described as transporter-related receptors and are present in many organisms (Thevelein & Voordeckers, 2009). Altogether, our data suggest that the role of CG1139 in infection may be signaling to orchestrate the responses necessary for the fast recovery of enterocytes. Moreover, gap junctions or connexin-like coupling was shown to be absent in the posterior midgut enterocytes. (S. R. Shanbhag, Vazhappilly, Sane, D’Silva, & Tripathi, 2017). Up to now, we have identified genes that are required for recovery in a non-cell autonomous manner, suggesting that cell to cell communication may be involved for the’ restoration of epithelial thickness.

The size of a given cell type in an organ is usually rather uniform, implying there are inherent processes that regulate the control of final cell size that remain poorly understood at present (Ginzberg, Kafri, & Kirschner, 2015). There are multiple mechanisms that contribute to the establishment of final cell size including nutrition, biogenesis, and most importantly cell division. Cytoplasmic extrusion as documented here provides a unique setting to study the regulation of cell size independently of most of the parameters cited above. Interestingly, we have recently established that the thickness of regrown enterocytes is slightly larger in enterocytes that have been exposed to *Sm*Db11hemolysin.

## Material and methods

### Fly husbandry

*Drosophila melanogaster* flies were raised at 25°C and nearly 60% humidity, 14h of daylight and fed a standard semi-solid cornmeal medium (6.4% (w/v) cornmeal (Moulin des Moines, France), 4.8% (w/v) granulated sugar (Erstein, France), 1.2% (w/v) yeast brewer’s dry powder (VWR, Belgium), 0.5% (w/v) agar (Sobigel, France), 0.004% (w/v) 4-hydroxybenzoate sodium salt (Merck, Germany)).

The wild-type flies used for experiments were white *wA5001* (Thibault et al., 2004) and the DrosDel *w^1118^* isogenic stock (*w^1118^ iso*) (Ryder et al., 2004). The following tissue-specific driver lines were used: enterocyte-specific driver line NP1, (also known as Myo31D), *NP-Gal4-tubGal80^ts^* (Cronin et al., 2009; Nehme et al., 2007); ubiquitous *Ubi-Gal4tubGal80^ts^*; fat body-specific *Yolk-Gal4*; enteroendocrine cells-specific *prospero-Gal4*; midgut progenitor-specific *esg-Gal4Gal80^ts^*; visceral muscles-specific *how>Gal4*; Malpighian tubules-specific *uro-Gal4* (X. Li, Rommelaere, Kondo, & Lemaitre, 2020). *CG1139* RNAi used for most of the experiments was the KK line from Vienna Drosophila RNAi Center (VDRC) (*CG1139* RNAi), and the same line isogenized to *w^1118^ iso* (CG1139 RNAi iso) as described in (Ferreira et al., 2014). RNAi lines for the different amino acid transporters were from VDRC. *InR* RNAi (#51518) and *Myc* RNAi (#51454) were obtained from the Bloomington Stock Center. The drivers were used either alone or crossed with *w^1118^ iso* as controls, or crossed with a RNAi or a reporter line. Crosses were performed at 25°C except for *Inr* and *Myc* which were performed at 18°C. The F1 progeny of these crosses was then placed at 29°C with 70% humidity for 6 days in order to induce the expression of the *Gal4tubGal80^ts^* transgenes. The *UAS-CG1139-GFP* was kindly provided by Prof. Deborah Goberdhan. For clonal analysis, the Flp-Out line w; *esgGal4tubGal80^ts^ UAS-GFP; UAS-flp Act>CD2>Gal4* from (Jiang et al., 2009) was used. *CG1139* knock-out mutant was generated by the Sino-French Hoffmann Institute CRISPR-Cas9 platform and comprises a 5bp deletion in the sequence causing a presumably truncated non-functional protein (**Supplementary Figure 3**). *CG1139* Knock-in mutant was generated by deleting and replacing the entire CDS region of CG1139 by cassette of Gal4 in Well Genetics (**Supplementary Figure 5**). Both mutants were isogenized to *w^1118^ iso*.

### Microbiology and Infections

*Serratia marcescens* strain Db11 (*Sm*Db11) was cultured on Lysogeny Broth (LB) agar plates with 100 μg/mL of streptomycin. The strain 21C4 *Sm*21C4 (Kurz et al., 2003) was cultured on LB-agar plates with 20 μg/mL chloramphenicol. The solid plates were placed overnight at 37°C to obtain colonies. For liquid cultures, one bacterial colony was taken off the solid plate and inoculated into 200 mL of liquid LB. These cultures were kept overnight at 37°C with agitation. All infections were performed using a final bacterial OD 600 (Optical Density at 600 nm) of 10, except for the survival where OD=1 was used. The OD was measured with a spectrophotometer and 50 mL of this solution was centrifuged at 4000 g/rcf for 10 minutes. The pellet was re-suspended in the appropriate volume of a 50 mM sucrose solution containing 10% of LB, in order to reach a final OD=10. 2 mL of this infection solution were added to two absorbent pads (Millipore AP1003700) that were placed at the bottom of medium-size vials (3.5 cm diameter). Twenty female flies of five to seven-day old were fed this infection solution, or sucrose 50 mM as a control at 29°C.

For **Figure 1A**, essential amino acids (MEM Amino Acids Solution 50X, Thermo Fischer Scientific #11130051) were added to the infection solution to a final 1x concentration. The 1x solution is composed of: L-arginine hydrochloride (126.4 mg/L), L-cysteine (24 mg/L), L-histidine hydrochloride (42 mg/L), L-isoleucine (52.4 mg/L), L-leucine (52.4 mg/L), L-lysine hydrochloride (72.5 mg/L), L-methionine (15.1 mg/L), L-phenylalanine (33 mg/L), L-threonine (47.6 mg/L), L-tryptophan (10.2 mg/L), L-tyrosine (36 mg/L) and L-valine (46.8 mg/L). Non-essential amino acids (MEM Non-Essential Amino Acids Solution 100X, Thermo Fischer Scientific #11140076) were added to a final 1x concentration. The 1x solution is composed of: glycine (15 mg/L), L-alanine (17.8 mg/L), L-asparagine (26.4 mg/L), L-aspartic acid (26.6 mg/L), L-glutamic acid (29.4 mg/L), L-proline (23 mg/L) and L-serine (21 mg/L).

### Fluorescent histochemical staining

Midguts were dissected in PBS and fixed for 30 minutes with 4% paraformaldehyde. Samples were washed three times with PBS-Triton X-100 0.1% (PBT 0.1%).

#### Actin staining

For actin staining midguts were incubated for 1h30 at room-temperature or overnight at 4°C in 10 μM Fluorescein Isothiocyanate (FITC) (Sigma-Aldrich #P5282) or Texas-Red labeled phalloidin (Invitrogen TM #T7471). Samples were then washed three times with PBT 0.1%.

#### GFP staining

To visualize the CG1139-GFP fusion protein, midguts were incubated with 1:500 anti-GFP (mouse) antibody (Roche #11814460001) for 3h. Samples were then washed three times with PBT 0.1% and incubated with a secondary goat anti-mouse FITC antibody (Abcam #6785) for 1h30. In the case of a co-staining with phalloidin and antibodies, the primary antibody was added alone for 3h, followed by 2h of incubation with a mix containing phalloidin and the secondary antibody. The visualization of CG1139 KI> UAS GFP was directly done without antibody staining.

#### Phospho 4EBP staining

Midguts were blocked 2h in PBT 0.1% with 2% bovine serum albumin (BSA). P-4EBP was detected with the anti-phospho-4EBP1 rabbit antibody from Cell Signaling (#2855). Midguts were incubated for 3h with 1:200 of the P-4EBP antibody, washed three times with PBT 0.1% and incubated with a secondary goat anti-rabbit FITC antibody (Abcam #6717).

All samples were mounted on diagnostic microscope slides (Thermo Fisher Scientific) with Vectashield plus DAPI (Vector Laboratories). Samples were observed using a LSM780 confocal microscope (Zeiss) or in Axioskop 2 microscope (Zeiss). All images were analyzed with the ImageJ/Fiji software.

### Classification of gut epithelium thickness

To determine the level of thinning and the recovery capacity, we used either qualitative or quantitative analysis. For qualitative analysis, midguts were classified into three different categories according to their epithelial thickness. Thick epithelium, which we represent in blue, corresponds to cells with the normal size, (around 20μm in length), with a clear dome shape characteristic of the intestine. Thin epithelium, which we represent in red, corresponds to cells that are very thin (around 5μm) and the normal dome shape of the cells is not observed. Semi-thin epithelium, which we represent in yellow, corresponds to cells with intermediate thickness (around 13μm), where the dome shape is still not completely defined. For quantitative analysis, pictures of midguts were acquired using Fluorescence Axioskope Zeiss and the length of gut enterocytes was measured using ImageJ/Fiji software. The measure for each midgut corresponds to the average of the measure between 10 enterocytes. Measures were taken every 5 to 10 enterocytes.

### RTqPCR

RNA was extracted from 5 or 10 midguts (without crop and Malpighian tubules) in triplicates. Midguts were crushed into 100 μL of TRI Reagent RT (Molecular Research Center) with 5% of bromoanisole (BAN, 98%, Molecular Research Center). Samples were vortexed, incubated 5 minutes at room temperature and centrifuged 10 minutes, 15 000 rcf/g at 4°C. The upper phase of the samples was collected, mixed with 350 μL of isopropanol and vortexed. Tubes were centrifuged at 18 000 rcf/g for 15 minutes at 4°C. The pellet was washed twice with 500 μL of ethanol 70% and dried. RNAs were then re-suspended in 25 μL of MilliQ water (Millipore). 1 μg of RNA was then used to generate cDNA by reverse transcription, using the iScript cDNA synthesis kit (Bio-Rad #1708890). The quantitative Polymerase Chain Reaction (qPCR) was performed with the Bio-Rad iQ TM SYBR Green Supermix kit and data were analyzed with the CFX384 system (Bio-Rad). mRNA quantitation was done by normalizing the amount of RNA detected for the gene of interest, with the control rp49 mRNA levels. The relative gene expression was calculated by normalizing the values obtained from sucrose fed flies (controls) versus challenged ones (infected with SmDb11). Primers used for rp49 were the forward 5’GACGCTTCAAGGGACAGTATCTG-3’ and the reverse 5’- AAACGCGGTTCTGCATGAG3’. Primers used for CG1139 were the forward 5’ACGTCAGCTTTTCGCAGGCCA-3’ and the reverse 5’- ACAACGCAGCCAGGGTGGAC-3’. Primers used for myc were the forward 5’ CAGTTCCAGTTCGCAGTCAA-3’ and the reverse 5’ AGATAAACGCTGCTGGAGGA-3’.

### Free amino acids quantification

Free amino acids were measured using the L-amino acid quantitation kit from Sigma-Aldrich (#MAK002). Females flies were either starved on sterile water, kept on their normal food, fed sucrose 50 mM or fed the usual infection solution for 3, 6 and 12h. The hemolymph from 20 flies or 10 midguts were collected into 10 μL of the kit buffer and processed according to the manufacturer’s instructions. Samples were run in biological duplicates.

### Click-it assay

The Click-it assay was performed with the Click-it AHA (L-azidohomoalanine) kit from Thermofisher (#C10102), the Alexa Fluor 488 alkyne dye (#A10267) and the Click-it Cell Reaction Buffer kit (#C10269). The reagents were prepared according to the manufacturer’s instructions. 50 μM of AHA diluted in PBS was injected in the hemolymph of female flies. Flies were placed on different conditions: i) their normal food ii) sterile water iii) sucrose 50 mM iv) *Sm*Db11 diluted in sucrose 50mM. Midguts were dissected in PBS 6h post-treatment, fixed for 20 minutes in 4% paraformaldehyde and washed with PBT 0.1%. To detect the injected AHA, midguts were stained for 30 minutes (protected from light) with 0.5 mL of the following mix: 437.5 μL of 1X Click-it Reaction Buffer, 10 μL of CuSO 4 , 50 μL of Click-it Buffer Additive and 2.5 μL of 1 mM Alexa Fluor 488 alkyne. Midguts were washed three times with PBT 0.1% for 15 minutes and mounted on microscopy slides as previously described. Samples were immediately observed at the confocal microscope and the fluorescence intensity was measured with the ImageJ software.

### Fitness parameters

The negative geotaxis assay was performed as described in (Linderman, Chambers, Gupta, & Schneider, 2012). Ten flies per vial were either infected with an OD600=10 of SmDb11 or fed sucrose as a control. After 3 or 16h of feeding, they were transferred in the vials designed for the assay without anesthesia. A line was drawn at 8 cm from the bottom of the tube. The number of flies crossing this line after 5 and 10 seconds was measured and plotted on a graph. Survival assays were performed using 20 flies per vial in triplicates. The experiments were conducted with five to seven-day old adult females at 29°C with 70% humidity. The flies were fed either sucrose 100 mM as a control or infected with an OD600=1 of SmDb11. Two absorbent pads were placed at the bottom of medium-size vials and 2 mL of sucrose or bacterial solution were added to the filters. Each day, flies that were alive were counted and 200 μL of 100 mM sucrose was added to the vials.

Bacterial loads in the midguts or crop were accessed 16h post *Sm*Db11 infection. Guts were dissected and pools of 3 midguts and crops were homogenized in 100μl of PBS1x and serial dilutions were performed to plate. Extracts were plated in LB agar containing 100 μg/mL of streptomycin and incubated at 37°C overnight to count total number of colonies.

Food intake was accessed either by quantifying the level of ingested blue dye or using the FLIC system. To access the level of ingested blue dye, flies were either fed in sucrose or *Sm*Db11, in a solution containing 5% of standard food blue dye. After 16h females were homogenized in 50μm of PBS1x and samples were centrifuged at maximum speed for 10min. Absorbance was accessed using the Varioscan and the wavelength was measured at 625 nm. The absorbance for each sample was calculated to the relative controls, where flies were subjected to the same conditions but without blue dye. Each sample corresponds to a pool of 10 females. To access the food intake using the FLIC system (Ro, Harvanek, & Pletcher, 2014) we placed one fly per well feeding either in sucrose or SmDb11 solution. 12 females were accessed per condition in each experimental replicate. Number of leaks was considered as a read-out for the food intake.

Defecation was accessed by feeding 10 females per vial either in sucrose or in *Sm*Db11, in a solution containing 5% of standard food blue dye. Fecal spots on the vial walls were counted.

### RNA sequencing

To analyze genes differentially regulated in NP> ctrl and NP> CG1139 RNAi flies after infections, flies were infected with SmDb11 or exposed to sucrose and midguts were dissected 3h, 8h, or 12h post infection. Each sample comprised 10 midguts (without crop and Malpighian tubules) and three replicate samples were generated per condition.

Samples were sent to BGI and RNA was sequenced using the DNBseq platform. We used the Agilent 2100 Bio analyzer (Agilent RNA 6000 Nano Kit) to perform the total RNA sample quality control RNA concentration, 28S/18S, and the fragment length distribution. Firstly, we removed the reads mapped to rRNAs and obtained raw data; then, we filtered out the low quality reads (More than 20% of the bases qualities are lower than 10), reads with adaptors and reads with unknown bases (N bases more than 5%) to get the clean reads. We assembled those clean reads into Unigenes, followed with Unigene functional annotation and calculated the Unigene expression levels and SNPs of each sample. Finally, we identify DEGs (differential expressed genes) between samples and performed clustering analysis and functional annotations. After filtering the reads, clean reads were mapped to reference genome using HISAT2 (Kim, Langmead, & Salzberg, 2015) and the average mapping ratio with reference genome was 94.08%. Clean reads were mapped to reference transcripts using Bowtie2 (Langmead & Salzberg, 2012), and gene expression level was calculated for each sample with RSEM (B. Li & Dewey, 2011). Based on the gene expression level, we used DEseq2 algorithms to identify the DEG (Differentially expression genes) between samples or groups. With DEGs we performed Gene Ontology (GO) classification, KEGG pathway classification and functional enrichment. We calculated false discovery rate (FDR) for each p-value, in general, the terms which FDR not larger than 0.01 are defined as significant enriched. We extracted here genes involved in cell growth where we identified genes involved in insulin signaling and the transcription factor myc.

The average mapping ratio with reference genome was 94.08%, the average mapping ratio with gene was 80.06% and a total of 17,003 genes were detected.

Gene Set Enrichment Analysis (GSEA) was performed as described (Subramanian et al., 2005) using GSEA v4.1.0. We used a gene set library from the FlyEnrichr (E. Y. Chen et al., 2013; Kuleshov et al., 2016) that distributes the genes according to biological processes from their Gene Ontology, “GO_Biological_Process_AutoRIF_Predicted_zscore”.

### Statistical analysis

Statistical analyses were performed using GraphPad software Prism 6 and R. For qualitative analysis of epithelium thickness, we used chi-square statistical tests. We also used Linear models (lm), or linear mixed-effect models (lmer) (Bates, Mächler, Bolker, & Walker, 2015)(lmer package lme4) if there were random factors. Significance of interactions between factors was tested by comparing models fitting the data with and without the interactions using analysis of variance (anova). Models were simplified when interactions were not significant. Pairwise comparisons of the estimates from fitted models were analyzed using lmerTest, lsmean, and multcomp packages.

## Acknowledgements

We thank Prof. Deborah Goberdhan to have kindly provided the UAS-CG1139-GFP line, the Vienna Drosophila Resource Center (VDRC, www.vdrc.at) and Bloomington Drosophila Stock Center (NIH P40OD018537) for their resource. We are grateful to Prof. Jiyong Lu and the SFHI CRISPR-Cas9 platform for generating the *CG1139* KO strain.

We thank Jenny Nguyen for expert technical help in some experiments and to Roenick Olmo for the help with the GSEA analysis.

This work has been funded by CNRS, University of Strasbourg, ANR grant ANR-16-CE13-0011-01 (ENTEROCYTE_PURGE_RECOVERY), Fondation pour la Recherche Médicale (Equipe FRM DEQ20090515394 to DF) and fellowship FDT20170437224 to CS. Our work is also partially sponsored by Infinitus, Inc (China). The funding sources had no role in the design of the study nor in its execution, analyses, interpretation of the data or decision to publish the results.

## Notes

### Competing Interest Statement

The authors have declared no competing interest.

