## Supplementary Material for "The amino acid transporter CG1139 is required for retrograde transport and fast recovery of gut enterocytes after *Serratia marcescens* intestinal infection"

**Supplementary Figure 1. Experimental design of the retrograde transport assay. (A)** Accurate quantification for **Figure 1B** in an independent experiment. Pictures of midguts were acquired for each condition and enterocyte thickness was measured using ImageJ software. Each dot represents the average of 10 independent measures in one midgut. Independent measures of enterocyte thickness for each midgut were taken every 5 to 10 enterocytes. Enterocytes are thin when infected at 3h and recover to the normal size in all the three solutions (*lm*, *** $p<0.001$). **(B)** The Click-it reaction is a non-radioactive technique that allows the detection of nascent proteins. AHA corresponds to a modified amino acid mimicking methionine that is normally incorporated into proteins. The detection is based on the reaction between an azide group on the modified amino acid and the alkyne group of a fluorophore. We have used the Thermofisher Click-it AHA kit in order to detect the presence of the modified amino-acid in the midgut after its injection in the hemolymph. The Click-it AHA kit was used to assess the possibility of a retrograde transport from hemolymph to the intestine. 50 µM of AHA were injected in the hemolymph of female flies. Flies were then placed on different conditions: food, sucrose 50 mM, water or infected with *Sm*Db11. Midguts were dissected 6h later, and stained with the alkyne probe to assess the fluorescence corresponding to the incorporated amino-acid.

**Supplementary Figure 2. Amino acid transporters involved in the fast recovery of the gut epithelium after *Sm*Db11 infection. (A)** Epithelium thickness 16h after *Sm*Db11 infection in flies that were knocked down for the different amino acid transporters in the gut enterocytes (with NP> driver). **(B)** Phylogenetic tree based on protein sequence from the different amino acid transporters used to screen in **(A).** Phylogenetic tree generated using UniProt. In red are shown the amino acid transporters that are required to contribute to gut recovery in **(A)**.

**Supplementary Figure 3. CG1139 is required for the fast recovery after infection. (A)** Efficiency of *CG1139* knockdown in the *NP*>*CG1139*-RNAi KK line from VDRC. *CG1139* expression was measured by RT-qPCR on 10 dissected midguts, in triplicates. Expression was determined relative to sucrose controls. *CG1139* was induced in the midgut during the recovery, 9h post-infection in control but not *NP*>*CG1139*-RNAi flies (one-way ANOVA, $p < 0.01$). **(B)** Epithelium thickness 16h after *Sm*Db11 infection or sucrose exposure (Not Infected) in *NP*>ctrl, *NP*>*CG1139*-RNAi KK line and *NP*>*CG1139*-RNAi GD line. **(C)** Representative map of the gene *CG1139* with respective exons in yellow. CRISPR knock-out mutant for *CG1139* comprises a deletion of 5bp in the sequence, causing a frame-shift mutation with a non-functional truncated protein. **(D)** *CG1139* expression level in control flies, *NP*> *CG1139* RNAi or in the CRISPR KO mutant 9 h post infection. Gene expression was measured by RT-qPCR on 5 dissected midguts, in triplicates. Expression of *CG1139* increases in control flies but not in *NP*> *CG1139* RNAi or in the CRISPR KO mutant (lmer, *** $= p < 0.001$).

**Supplementary Figure 4. Knockdown of CG1139 in other host tissues and cell types (A).** *CG1139* was knocked down in enterocytes (*NP*> *CG1139* RNAi), whole body (*ubi*> *CG1139* RNAi) and in the fat body (*NP*> *CG1139* RNAi). Flies were infected with *Sm*Db11 and thickness of gut epithelium was assessed 16h post infection. Knocking down *CG1139* in whole body and in the enterocytes delays the recovery to the thick gut epithelium. **(B)** *CG1139* was knocked down in enteroendocrine cells (*prospero*> *CG1139* RNAi *iso*). Flies were infected with *Sm*Db11 or exposed to sucrose and thickness of gut epithelium was assessed 3h and 16h post infection. **(C)** *CG1139* was knocked down in the visceral muscle (*how*> *CG1139* RNAi *iso*) and in the Malpighian tubules (*uro*> *CG1139* RNAi *iso*). Flies were infected with *Sm*Db11 or exposed to sucrose and thickness of gut epithelium was accessed 16h post infection. **(D)** *CG1139* was knocked down in the enterocytes (*NP*> *CG1139* RNAi iso) or in the intestinal stem cells (*esg*> *CG1139* RNAi *iso*) and flies were exposed to sucrose for 16h. This is the control for **Figure 2H**.

**Supplementary Figure 5 – Midgut cells expressing CG1139. (A)** Scheme for the Knock-in CRISPR mutant generated for *CG1139* (*CG1139* KI CRISPR mutant). Entire coding sequence of *CG1139* was deleted and replaced by a Gal4 cassette. **(B)** *CG1139* KI CRISPR mutant was crossed with UAS-GFP and these flies were exposed to sucrose or infected with *Sm*Db11 for 8h and 20h. Cells expressing *CG1139* are GFP positive cells (green). Blue = DAPI; Red = Actin. Expression of *CG1139* is present in progenitor cells after sucrose exposure and also after infection. Expression of *CG1139* is also observed in some enterocytes. *CG1139* is also expressed in Malpighian tubules and posterior midgut, either in sucrose and infection conditions.

**Supplementary Figure 6 – Knockdown of *CG1139* in the gut enterocytes has an impact on *Sm*Db11 bacterial loads in the gut upon infection. (A)** Fly locomotion accessed by Negative Geotaxis Assay in control and *NP*> *CG1139* RNAi flies. Flies were either exposed to sucrose or infected with *Sm*Db11 and the assay was performed 3h and 16h post-infection. There are no significant differences between control and *NP*> *CG1139* RNAi flies (One-way ANOVA) **(B)** Survival of *NP*>ctrl and *NP*>*CG1139*-RNAi flies during chronic infection with *Sm*Db11 OD=1 or exposed to sucrose 50mM. There is no difference in survival between *NP*> ctrl and *NP*> *CG1139* RNAi flies (*lmer*, *p*-value>0.05) **(C)** Bacterial loads 16h after *Sm*Db11 infection in *NP*>ctrl or *NP*>CG1139-RNAi. Each sample (each dot) corresponds to a pool of three midguts or three crops. *NP*> *CG1139* RNAi have higher bacterial loads in the midgut and crop compared to control flies (*lmer*, ***=*p*<0.001). **(D)** *NP*> ctrl and *NP*> *CG1139*

RNAi flies were either infected or exposed to sucrose in a solution containing blue dye for 16h. Flies were collected, and food intake was analyzed by measuring the absorbance of blue dye ingested in the Varioscan. **(E)** *NP*> ctrl and *NP*> *CG1139* RNAi flies were infected or exposed to sucrose using the FLIC system for 16h and number of events was quantified. **(F)** *NP*> ctrl and *NP*> *CG1139* RNAi flies were either infected or exposed to sucrose in a solution containing blue dye for 16h. Fecal spots were counted in the tube. There is no significant difference between the fly lines (*lm, p*>0.05).

**Supplementary Figure 7. Categories of genes significantly enriched in *NP*> ctrl and not in *NP*> *CG1139* RNAi upon infection.** Gene set enrichment analysis was performed in the RNA sequencing results to compare enriched gene sets between *NP*> ctrl and *NP*> *CG1139* RNAi. **(A)** Genes involved in positive regulation of growth, lamellipodium formation and positive regulation of translation are positively enriched at 8h post infection in *NP*> ctrl and not in *NP*> *CG1139* RNAi. **(B)** At 16h post infection, there is an enrichment of genes involved in lamellipodium formation, actin nucleation and negative regulation of growth in the *NP*> ctrl and not in *NP*> *CG1139* RNAi.

**Supplementary Figure 8. Genes involved in growth differentially expressed in *NP*> *CG1139* RNAi upon infection. (A)** Genes involved in growth that are up- or down-regulated in *CG1139* RNAi 3h and 8h after infection (relative to sucrose). **(B)** Expression of *Myc* was measured by RT-qPCR in dissected midguts of *NPiso*> *CG1139* RNAi and $w^{1118}$ *iso*> *CG1139* RNAi flies 8h and 16h post infection. *Myc* is down-regulated at 8h in both flies but at 16h is only down-regulated in *NPiso*> *CG1139* RNAi, and not significantly different in control flies (*lmer*, p = 0.0116).

**Supplementary Figure 9. TOR pathway activity decreases in CG1139-RNAi flies. (A)** Confocal pictures of dissected midguts. Blue=DNA ; Green=phospho-4EBP (P4EBP). Midguts were stained with the P-4EBP antibody to detect when and where the TOR pathway was active. P-4EBP signal decreased 16h post-infection in *CG1139* knock-down flies, as well as when fed sucrose for 16h.

#### Supplementary Figure 1

A

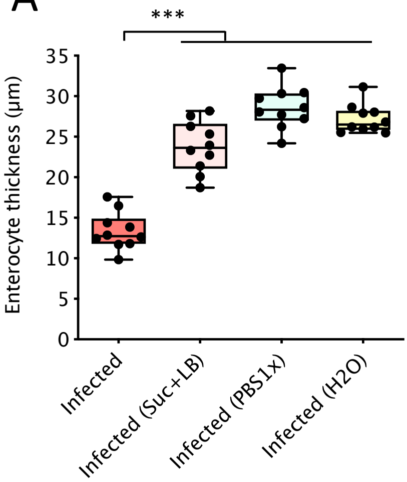


B

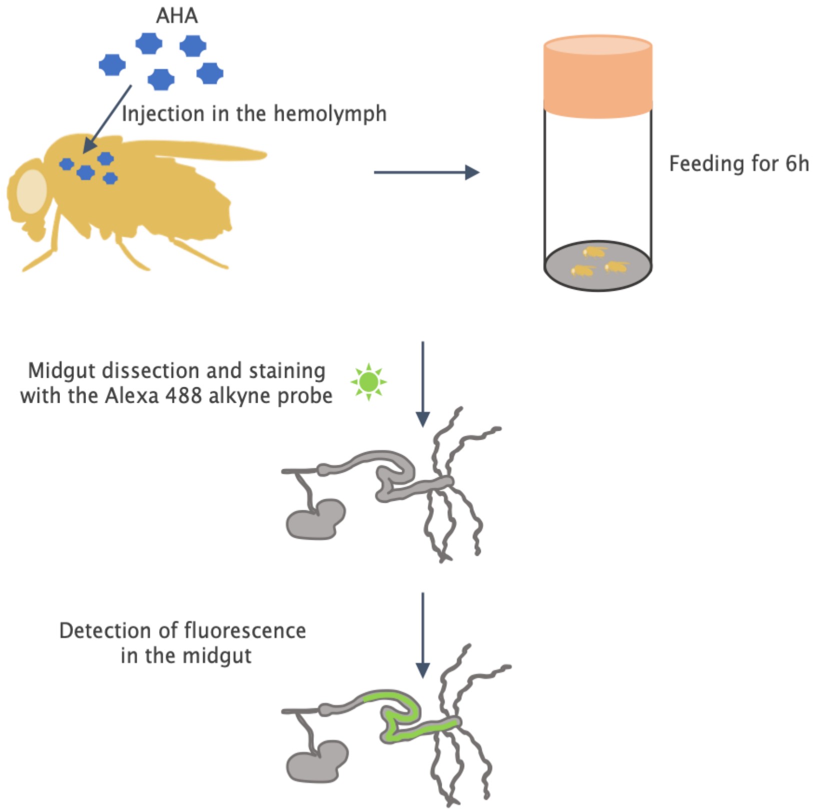

#### Supplementary Figure 2

A

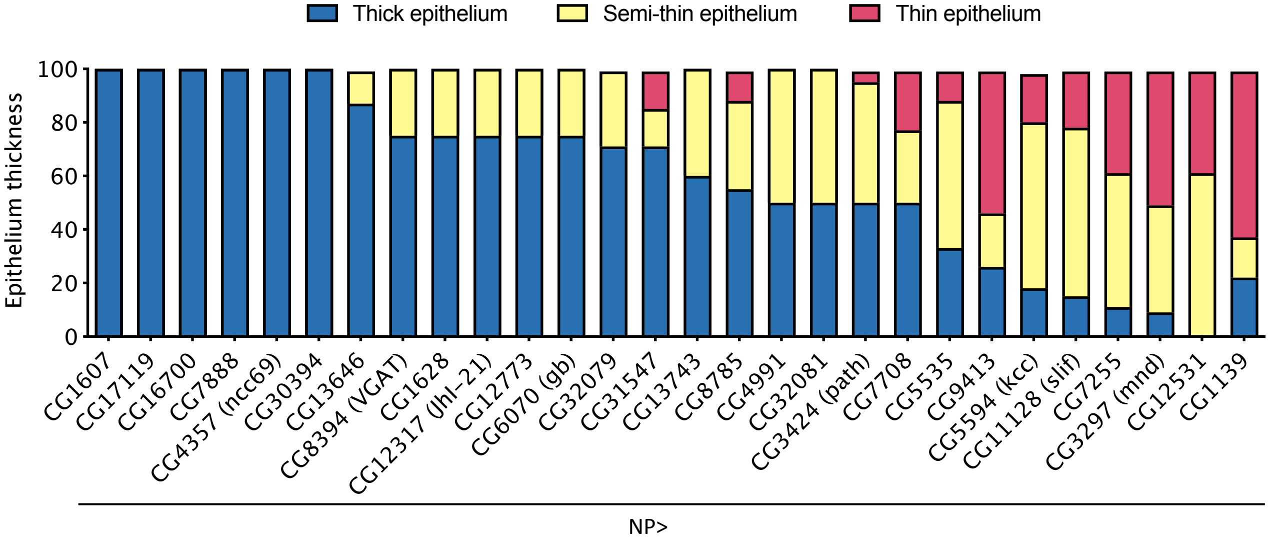


B

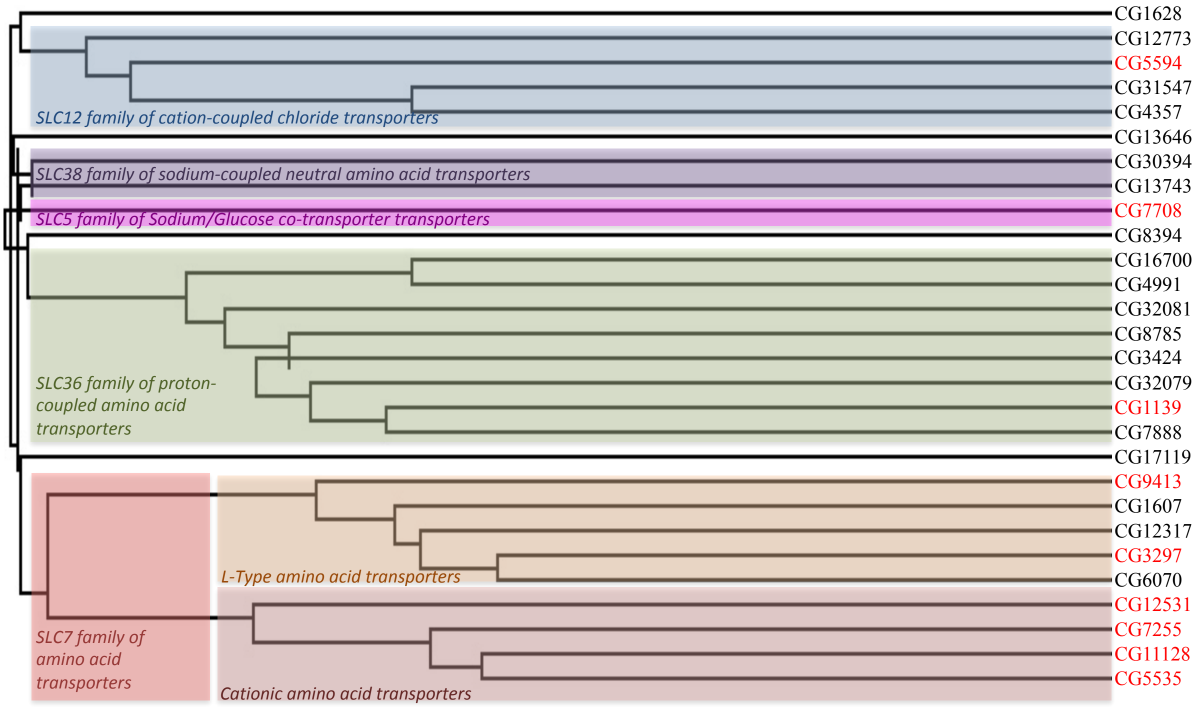

#### Supplementary Figure 3

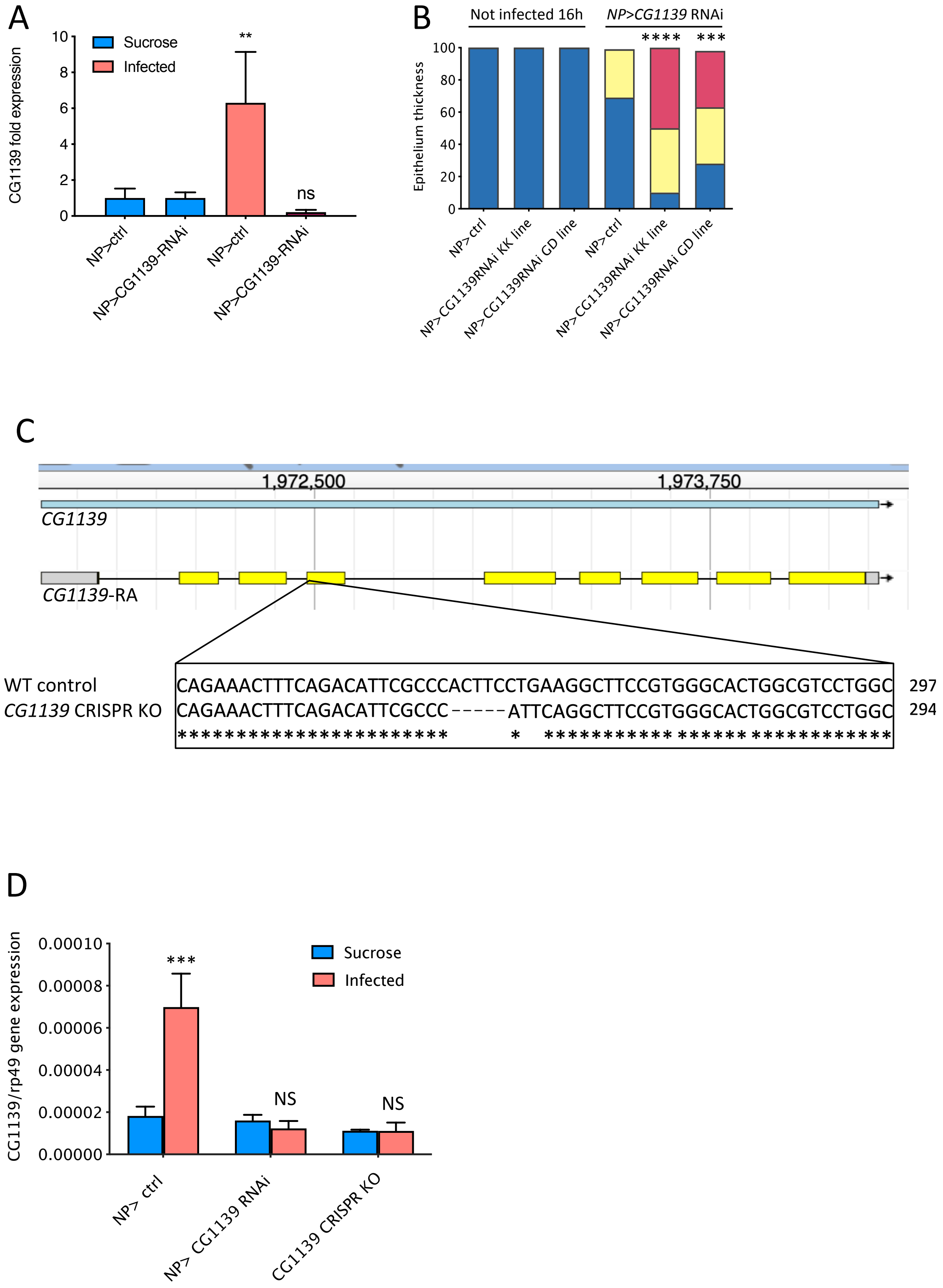

### Supplementary Figure 4

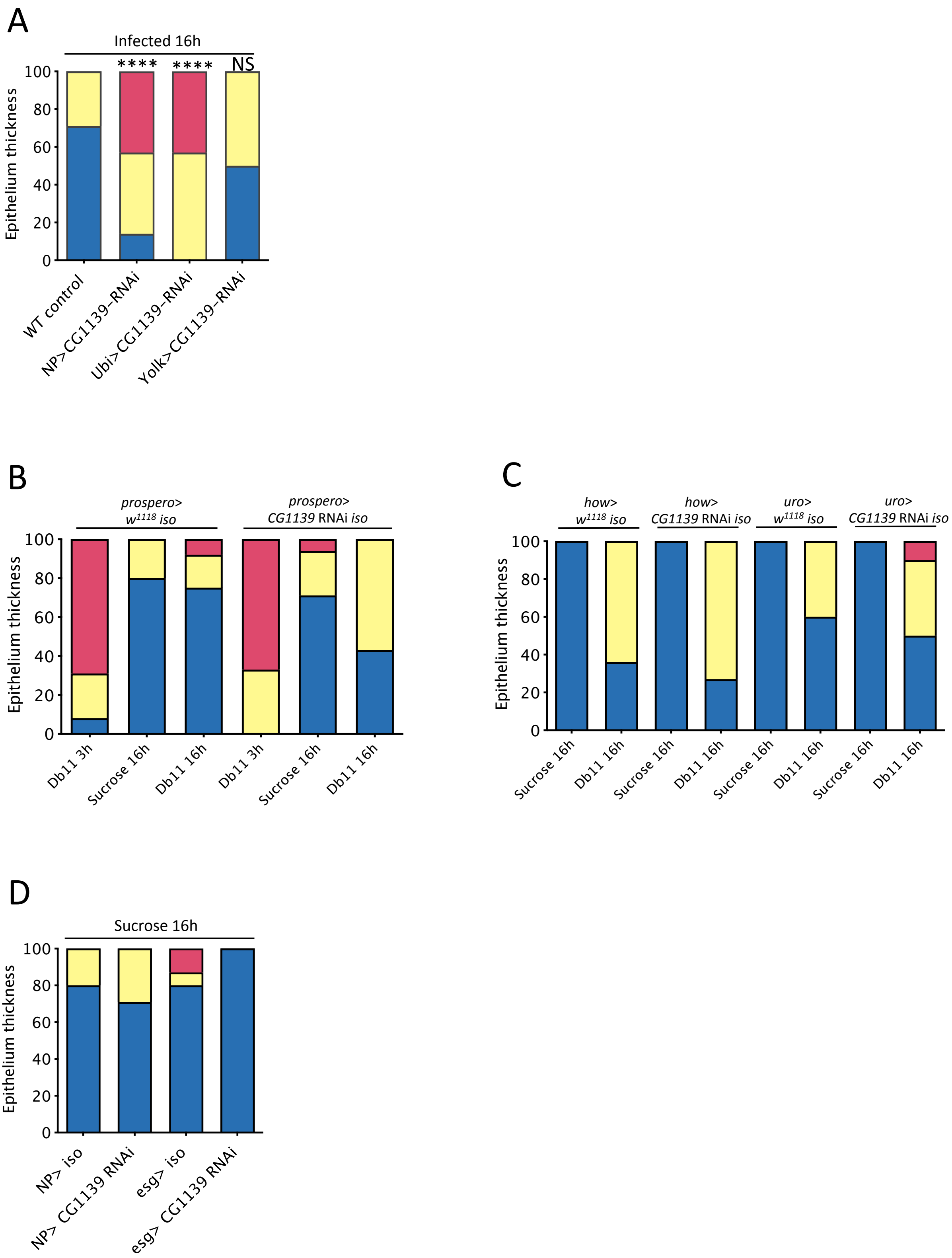

#### Supplementary Figure 5

A

**Injection strain:** *w[1118]*

**Genome Editing:**

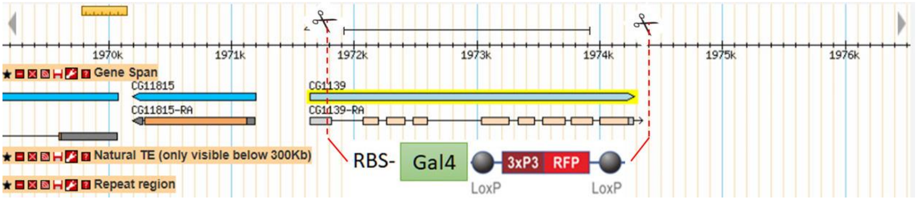


(1) The entire CDS region of *CG1139* will be deleted and replaced by cassette Gal4-RFP.

(2) Cassette Gal4-RFP contains ribosome binding sequence (RBS), Gal4, SV40 polyA terminator, and floxed 3XP3-RFP as a selection marker.

(3) The selection marker 3XP3-RFP contains loxP site, 3X Pax3 promoter, RFP, alpha-Tubulin 3'UTR and loxP site. It facilitates the genetic screening and can be flipped out by Cre recombinase.

B

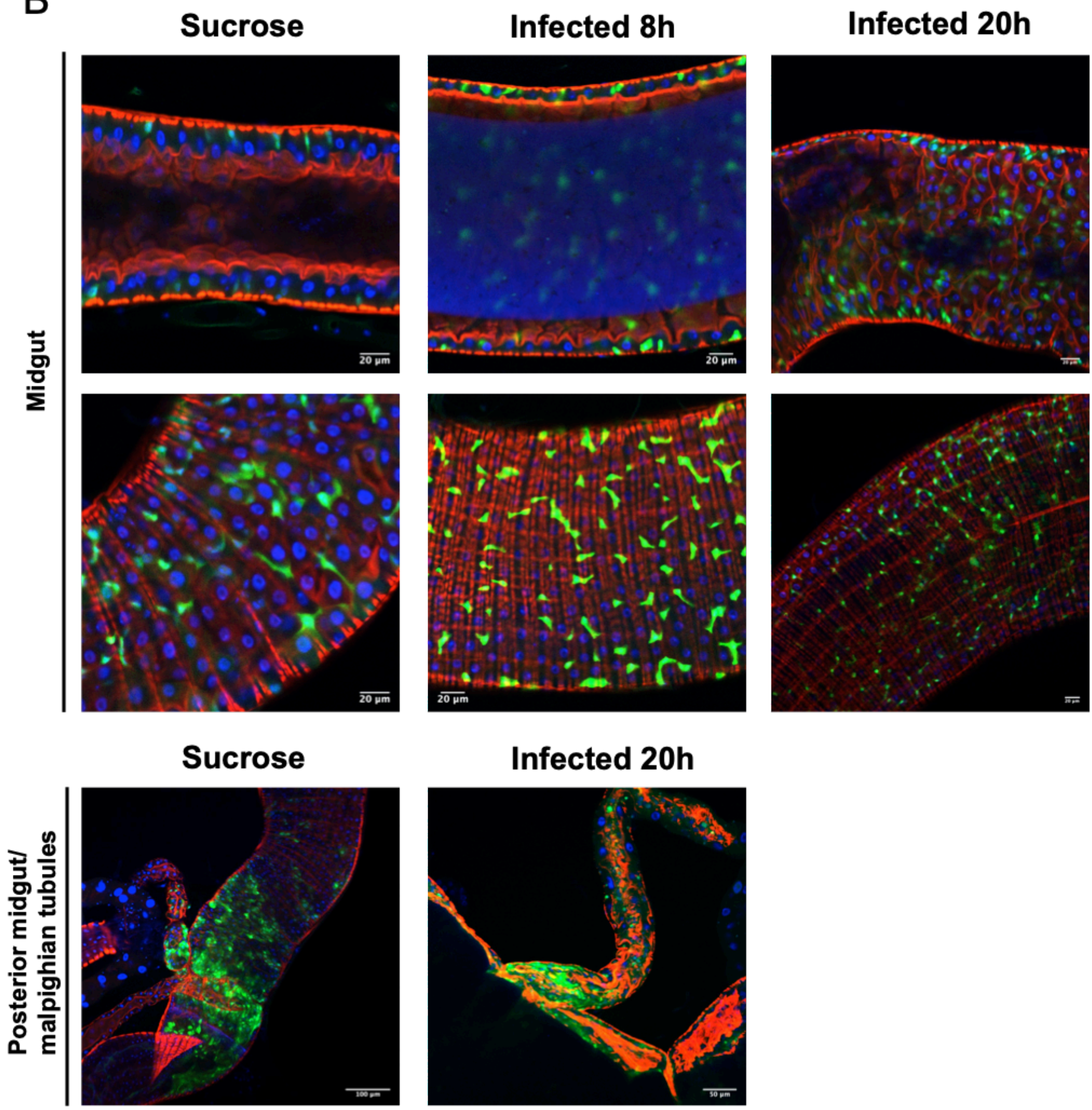

#### Supplementary Figure 6

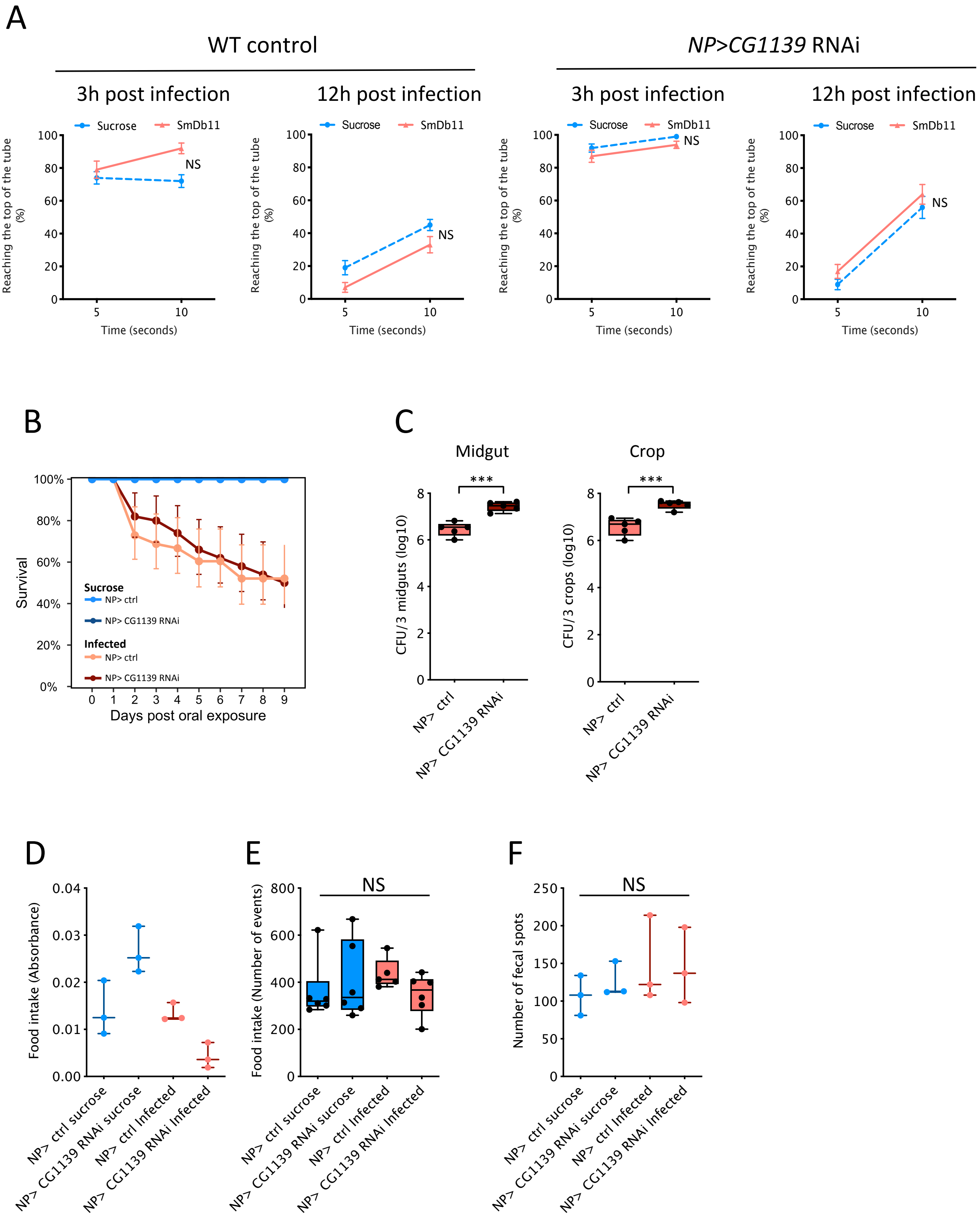

### Supplementary Figure 7

## A

**Gene sets positively enriched in the control and not in *NP> CG1139* RNAi at 8h**

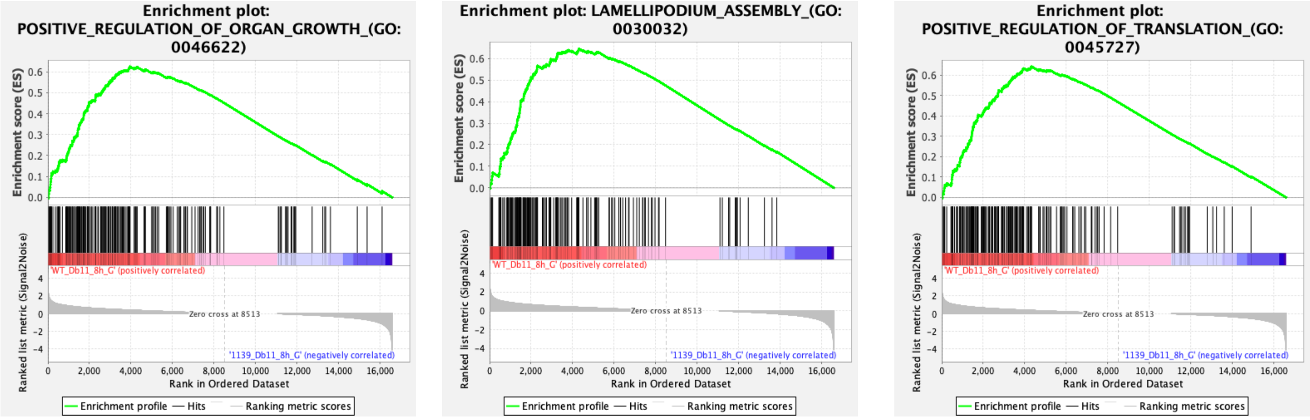


## B

**Gene sets positively enriched in the control and not in *NP> CG1139* RNAi at 16h**

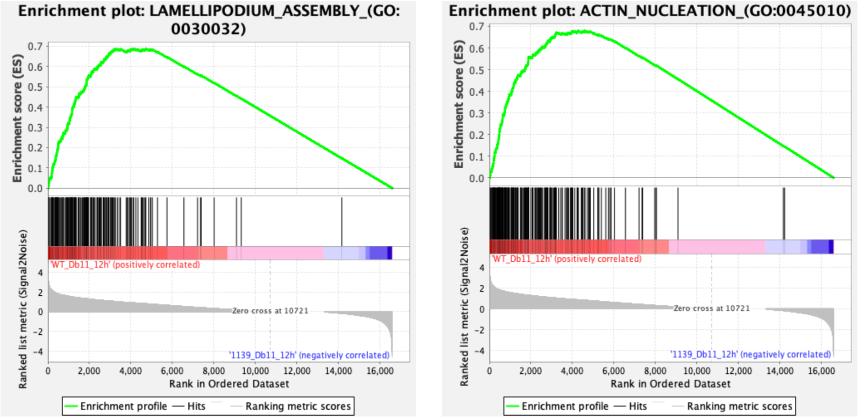

### Supplementary Figure 8

A

**Genes involved in growth differentially expressed in *NP> CG1139* RNAi (Infected/Sucrose)**

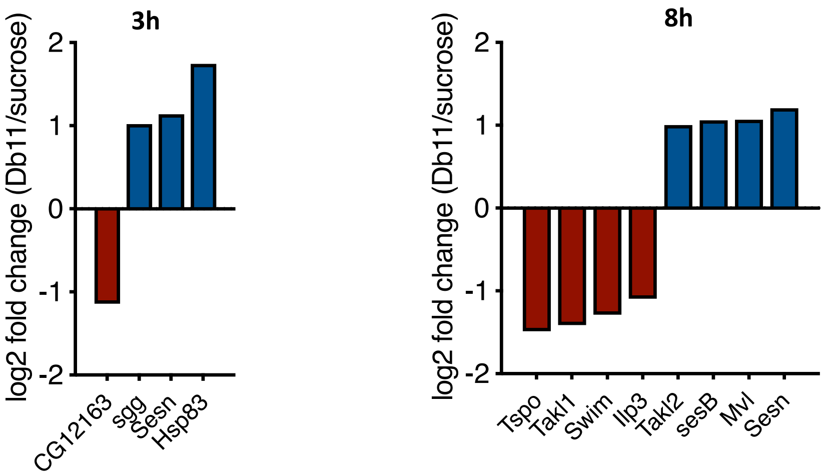


B

***Myc* expression levels in the gut after *Sm*Db11 infection**

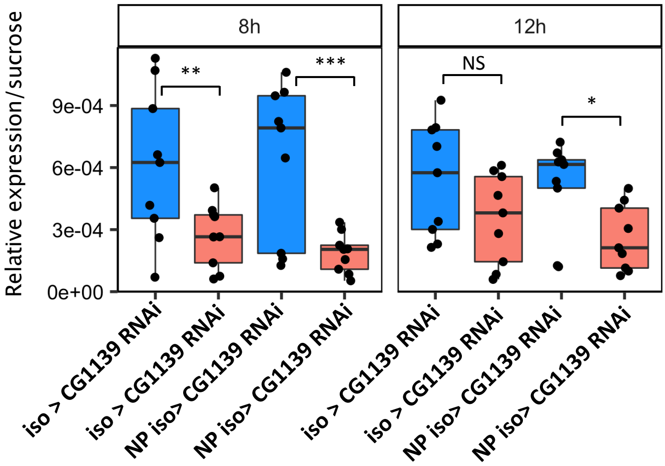

**Supplementary Figure 9**

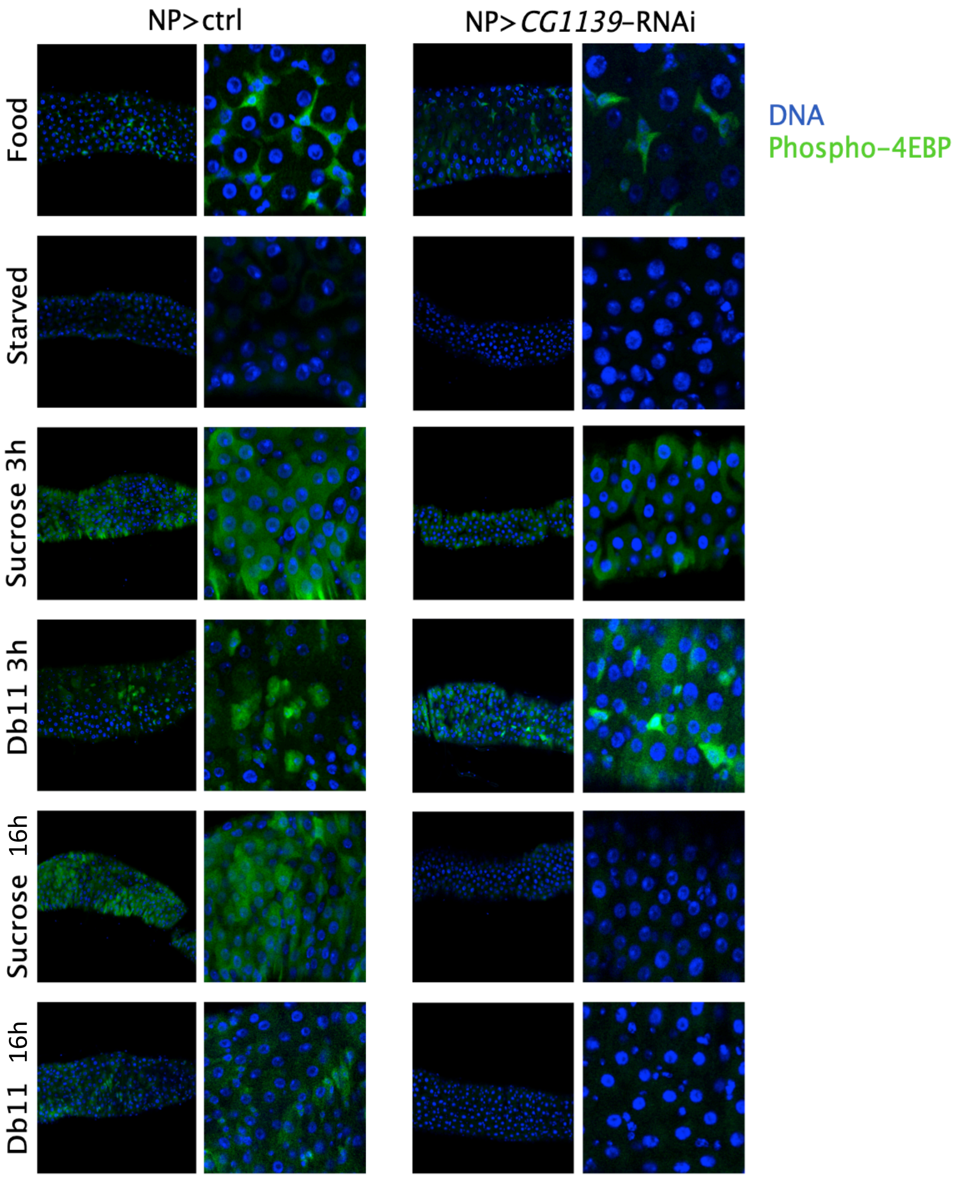
